# Functional hotspots identification *via* a hybrid NMR-computational approach facilitating directed evolution of large enzyme

**DOI:** 10.1101/2025.03.12.642771

**Authors:** Yihao Chen, Zhou Gong, Xiaoling Zhao, Ali Raza, Mingjun Zhu, Zhiqing Tao, Wenhui Li, Xu Zhang, Maili Liu, Lichun He

## Abstract

Directed evolution has revolutionized enzyme engineering but remains challenging for large, multidomain proteins due to their complex dynamics and vast combinatorial sequence space. Traditional NMR spectroscopy, while powerful for identifying functional residues, is limited by the requirement for complete resonance assignments that become intractable in high molecular weight systems. Here, we present a hybrid NMR-computational framework that integrates chemical shift perturbation mapping with AlphaFold3-docked distance constraints, enabling the identification and assignment of ligand-proximal hotspots without the need for full resonance assignment. By analyzing side chain ^1^H-^13^C correlation spectra, probabilistic amino acid typing, and structural proximity filtering, we mapped functional hotspots in the ∼90 kDa *Pyrococcus furiosus* DNA polymerase. Targeted saturation mutagenesis of these sites resulted in variants exhibiting multi-fold increases in catalytic efficiency while sampling less than 0.6% of total residues. The identified hotspots revealed spatially distinct yet cooperative contributions to catalysis, as further supported by combinatorial mutagenesis and kinetic analyses. This hybrid strategy provides a practical pipeline for narrowing mutagenesis targets in large enzymes under catalytic conditions, overcoming the size, assignment, and screening limitations of conventional NMR-guided evolution, offering a powerful route toward efficient biocatalyst optimization by prioritizing high-probability target residues.

## Introduction

Native enzymes, refined over hundreds of millions of years of evolution, possess functional cores optimized for organismal survival, however, they are not necessarily optimized for maximal catalysis of specific chemical reactions under industrial conditions (Sheldon and Woodley, 2018; Lovelock *et al*, 2022; Lister *et al*, 2025; Shi *et al*, 2023). Consequently, researchers are no longer limited to discovering natural enzymes but are actively engaged in engineering or designing enzymes tailored for specific applications (Arnold, 2018). To this end, directed evolution has emerged as a powerful strategy to address key limitations such as insufficient catalytic efficiency, narrow substrate scope, and poor stability under non-physiological conditions by enhancing critical biocatalyst properties including stability, activity, and selectivity (Forget *et al*, 2025; Sun *et al*, 2022; Bell *et al*, 2022; Zhang *et al*, 2022; Carr *et al*, 2003). These advances have facilitated the broad application of engineered enzymes, thereby revolutionizing fields such as biotechnology and biomedical research (Doudna and Charpentier, 2014; Turner, 2009; Hafeez *et al*, 2020). Developing broadly applicable methodologies for directed enzyme engineering is therefore a significant pursuit. However, evolving native enzymes remains challenging due to their high molecular weight, multi-domain architectures, and complex substrates (Soskine and Tawfik, 2010; Geddie and Matsumura, 2004). Even when the substrate binding pocket is known, it still takes a lot of time to mutagenize all the residues that cover the pocket, not to mention the combinatorial explosion that occurs (Wang *et al*, 2021). Moreover, enzymes depend on dynamic residue interactions beyond static active sites. These features avoid conventional structural analysis (Otten *et al*, 2020; Acevedo-Rocha *et al*, 2021; Bhattacharya *et al*, 2022; Gutierrez-Rus *et al*, 2025; Marshall *et al*, 2023). Therefore, without high throughput screening capabilities, experimental interrogation of exhaustive mutant libraries demands prohibitive resources, particularly when addressing the combinatorial complexity of large enzymes (Zhang *et al*, 2022; Marshall *et al*, 2023; Yu *et al*, 2022). Therefore, narrowing sequence search spaces is supreme, driving substantial efforts to develop methods for designing focused libraries to guide directed evolution (Bhattacharya *et al*, 2022; Gutierrez-Rus *et al*, 2025; Marshall *et al*, 2023; Khersonsky *et al*, 2018; Wijma *et al*, 2015).

In this context, nuclear magnetic resonance (NMR) spectroscopy provides a complementary strategy by enabling direct analysis of catalytic parameters and residue specific chemical environments under catalytic conditions (Otten et al, 2020; Bhattacharya *et al*, 2022; Hu *et al*, 2021). By exploring enzymes in their apo- and holo- states in solution, NMR can capture functional hotspots, thereby focusing evolutionary efforts on catalytic relevant residues and facilitating directed evolution of enzymes (Bhattacharya *et al*, 2022; Gutierrez-Rus *et al*, 2025). Moreover, NMR spectroscopy could also resolve residue specific chemical environments of enzymes under catalytically relevant conditions with cofactors (Mainz *et al*, 2013). In particular, 2D [^13^C,^1^H]-Heteronuclear Multi-Quantum Coherence (HMQC) experiments offer a powerful tool to probe the chemical environment of hydrophobic cores and functional sites (Tugarinov *et al*, 2007; Ruschak and Kay, 2010; Korzhnev *et al*, 2004) making them particularly suitable for capturing key functional hotspots in directed evolution studies. This is because the methyl groups of hydrophobic residues are highly sensitive to local conformational changes and can reveal subtle perturbations in protein dynamics or allosteric networks, even in large molecular systems. Nevertheless, the assignment of these hotspots is challenging, especially for large enzymes, where fast transverse relaxation rates make scalar magnetization transfer inefficient (Frueh, 2014). Consequently, there is an urgent need for more effective assignment methods to facilitate NMR analysis of hotspots of large enzymes.

Here, we developed a hybrid NMR-computational approach for mapping functional hotspots and guiding directed evolution of large enzymes. This method integrates NMR derived chemical shift perturbations (CSPs) with computationally derived distance constraints. By matching CSPs to these distance constraints and deriving structural insights, it predicts and assigns key functional hotspots, providing an alternative while complete NMR assignment is impractical. The high-fidelity DNA polymerase from the thermophilic archaeon *Pyrococcus furiosus* (PfuPol) with molecular weight ∼90 kDa was applied as a model system (Lundberg *et al*, 1991). Hotspots across spatially distinct regions of PfuPol were identified, effectively narrowing sequence search space. Saturation mutagenesis of identified hotspots yielded substantial improvements in separate biocatalytic functions, demonstrating the feasibility and effectiveness of the integrative approach in directed evolution of large enzymes, paving the way for new fundamental studies on enzyme functions and evolution.

## Results

### Hybrid NMR-computational strategy and workflow

NMR directed evolution of proteins including enzymes usually relied on the complete assignment of the target protein. However, for large enzyme, the complete assignment is hard to achieve. Therefore, to enable the NMR aided identification of functional hotspot, we developed a hybrid NMR-computational approach encompassing five sequential steps: (a) Determination of catalytic hotspots *via* NMR spectroscopy derived ^1^H-^13^C chemical shift perturbation of target enzymes upon adding substrates, neutralization antibodies or cofactors (Figure 1A); (b) Computational prediction of amino acid types based on Biological Magnetic Resonance Bank (BMRB) database (Figure 1B); (c) Integrated interpretation of CSPs with AlphaFold3-docked distance constraints to assign the hotspots. (Figure 1C); (d) Site-directed saturation mutagenesis of assigned hot spots for screening (Figure 1D); and (e) Exploration of combinatorial mutations coupled with functional characterization (Figure 1E). We applied this strategy to PfuPol which is central to molecular biology technologies, driving the translation of theoretical insights into applications (Moore, 2005; Shendure *et al*, 2017; Gibson *et al*, 2009).

**Figure 1.**
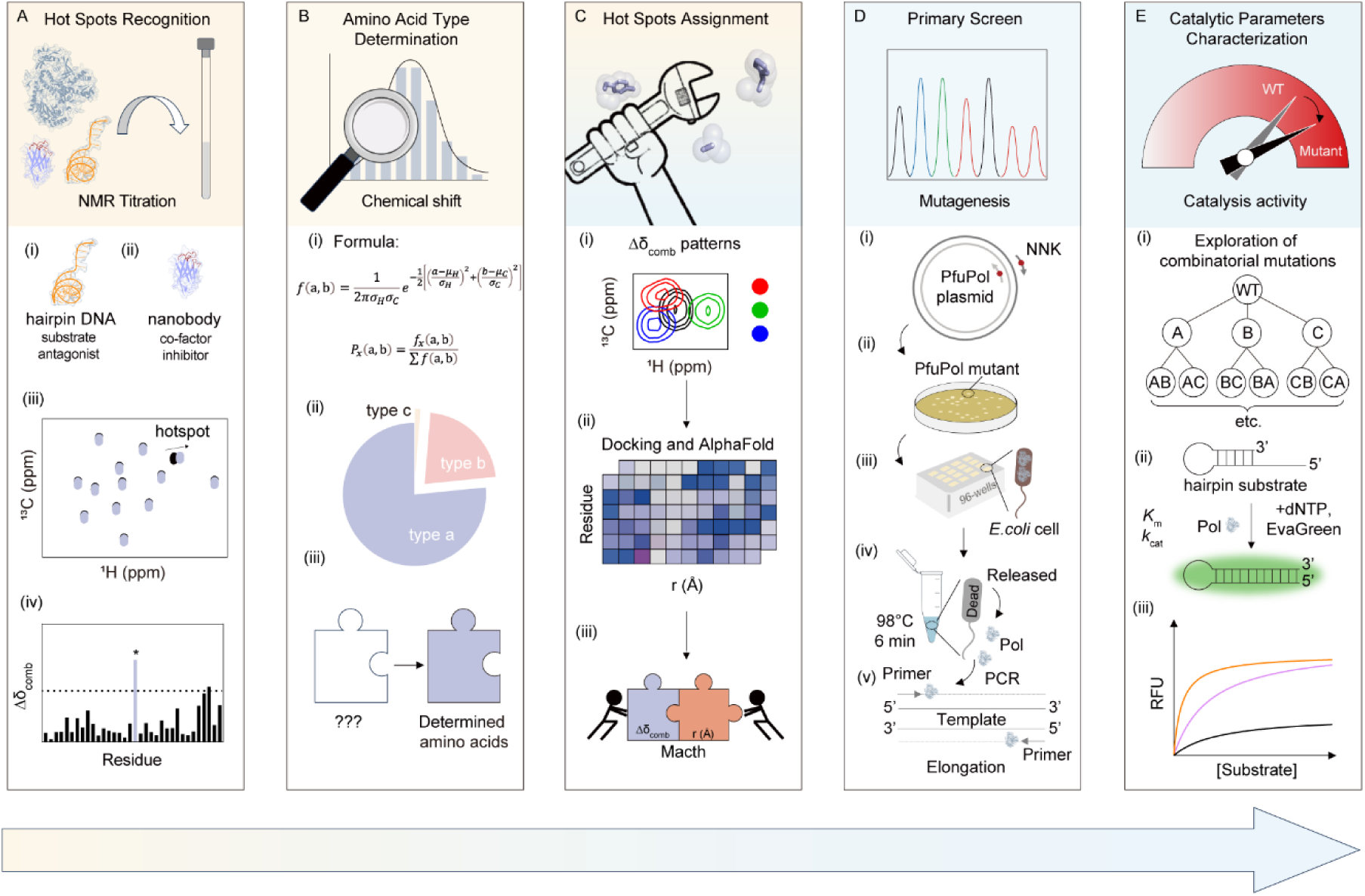
Schematic series of the hybrid NMR-computational approach: from functional hotspot mapping to guided evolution of large enzymes. (**A**) Schematic representation of hotspots identification by NMR titration. (**B**) Schematic diagram of the determination of amino acid types of hotspot residues. (**C**) Assignments of hotspot residues. (**D**) Schematic pipeline for the primary screening of hotspot residue libraries. (**E**) Schematic workflow for characterization of catalytic parameters and exploration of combinatorial mutations.

### Identification of hotspots

To enable NMR chemical shift perturbation experiments, we developed a hairpin DNA substrate (Moody *et al*, 2023) for NMR titration with PfuPol (Figure 1A(i) and Supplementary Figure S2A). Notably, we observed that the hairpin DNA exhibits a concentration dependent inhibitory effect in conventional PCR reactions (Supplementary Figure S2B). Therefore, we conclude that at micromolar concentrations, this hairpin DNA effectively occupies the active site of PfuPol in our NMR chemical shift perturbation experiments, thereby facilitating the identification of its functional hotspots. As shown in Figure 1A(iii) and (iv), we quantified CSPs (Δδ_comb_) by calculating standard deviation from the differences in the ^13^C and ^1^H chemical shifts of the PfuPol upon binding the substrate analogue. Following the established protocol, 2D [^13^C,^1^H]- Heteronuclear Multi-Quantum Coherence (HMQC) spectra of [^13^C]-labeled PfuPol were recorded both in the absence and presence of the hairpin DNA template (Figure 2A and Supplementary Figure S3A). Significant CSPs were observed for one aliphatic residue (‘ali1’) and three aromatic residues (‘aro1’, ‘aro2’, ‘aro3’) (Yu *et al*, 2017; Aguirre *et al*, 2014), with CSP magnitudes exceeding three times the standard deviation (>3σ) (Figure 2D).

**Figure 2.**
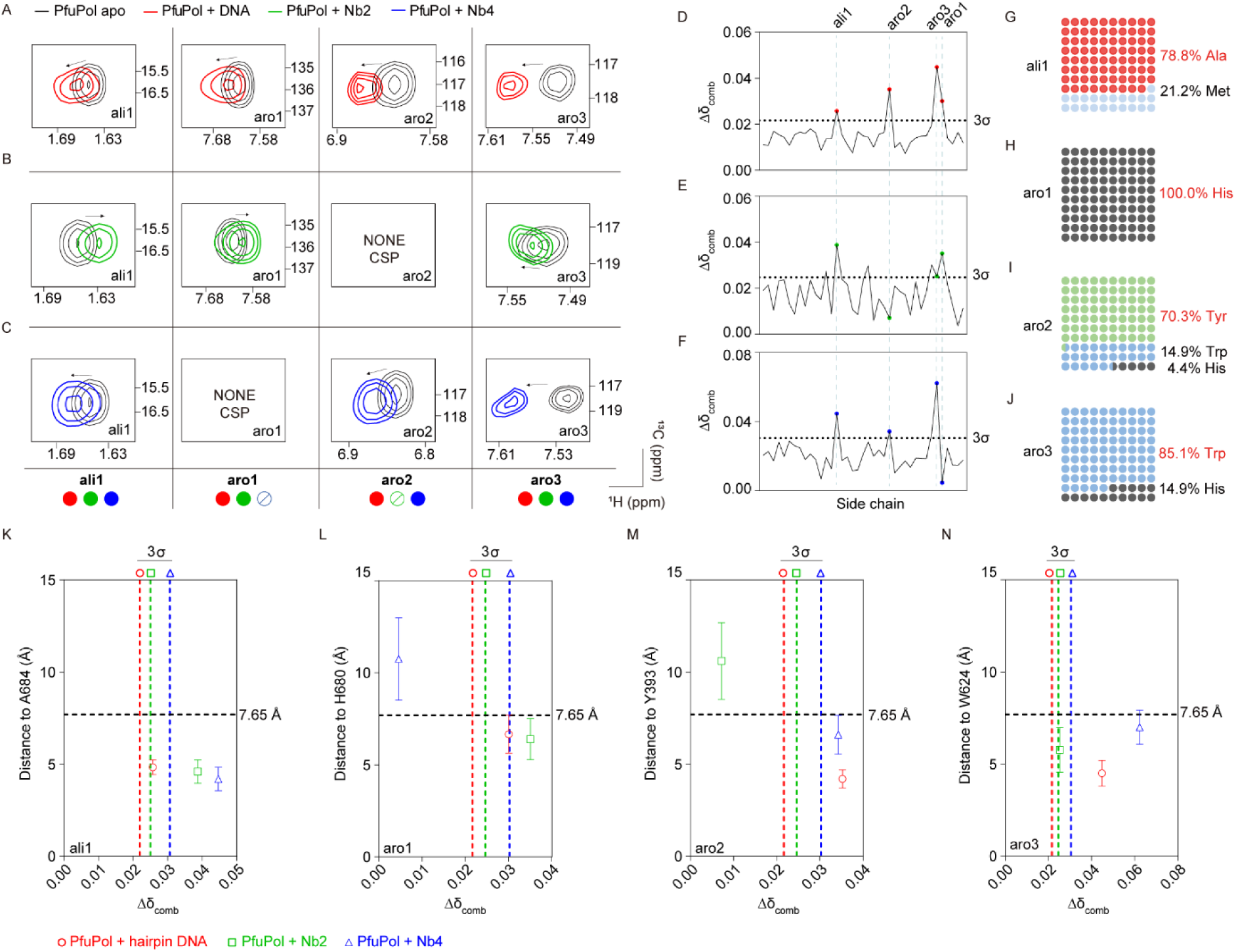
Identification and assignment of hotspots of PfuPol. (**A–C**) Two-dimensional [^13^C,^1^H]-HMQC spectra of 50 μM [^13^C]-labeled PfuPol were acquired on the 600 MHz spectrometer at 298 K. Enlarged view of NMR peaks of hotspot residues, which show CSPs upon adding 50 μM hairpin DNA (**A**, red), 75 μM Nb2 (**B**, green), 75 μM Nb4 (**C**, blue). Residues with CSPs were depicted by a colored circle at the bottom. Red circle indicated hairpin DNA induced CSPs. Green circle indicated Nb2 induced CSPs. Blue circle indicated Nb4 induced CSPs. Circle with a slash denoted no CSPs. (**D**) Side chain CSP of PfuPol upon addition of 50 μM hairpin DNA. (**E**) Side chain CSP of PfuPol upon addition of 75 μM Nb2. (**F**) Side chain CSP of PfuPol upon addition of 75 μM Nb4. (G–J) The probability of each amino acid type for residues ali1 (**G**), aro1 (**H**), aro2 (**I**), and aro3 (**J**) was determined based on calculations using probability density functions and Bayes’ theorem. (**K–L**) The minimum distances (r) between residues in PfuPol and their interaction partners (DNA, Nb2, Nb4) were plotted alongside the chemical shift perturbations (Δδ_comb_) observed at the hotspots upon addition of the interaction partners. (**K**) The assignment of ali1 was made to residue A684, which exhibited a minimum distance of less than 7.65 Å from all three binders (hairpin DNA, Nb2, and Nb4) and showed significant perturbation upon interaction with each of them. (**L**) The assignment of aro1 was made to residue H680, which exhibited a minimum distance of less than 7.65 Å with each of hairpin DNA and Nb2 and showed significant perturbation upon interaction with each of them. (**M**) The assignment of aro2 was made to residue Y393, which exhibited a minimum distance of less than 7.65 Å with each of hairpin DNA and Nb4 and showed significant perturbation upon interaction with each of them. (**N**) The assignment of aro3 was made to residue W624, which exhibited a minimum distance of less than 7.65 Å from all three binders (hairpin DNA, Nb2, and Nb4) and showed significant perturbation upon interaction with each of them.

As the chemical shift perturbation may arise from the binding sites and the allosteric conformational changed regions which may not directly involved in the catalytic activity of PfuPol, two neutralizing nanobodies (Nb2/Nb4) derived from our prior work (Tao *et al*, 2024) were then employed in NMR titration to filter non-catalytically relevant hotspot (Figure 1A(ii) and Supplementary Figure S2C–E). Similarly, 2D [^13^C,^1^H]-HMQC spectra of [^13^C]-labeled PfuPol were recorded in the absence and presence of unlabeled nanobodies Nb2 and Nb4, respectively (Supplementary Figure S3B and C). The presence of the nanobody Nb2 induced notable CSPs (>3σ) of peaks ali1, aro1, and aro3 (Figure 2B and E), while the presence of Nb4 caused pronounced significant CSPs (>3σ) of peaks ali1, aro2, and aro3 (Figure 2C and F). Residues ali1, aro1, aro2, and aro3 revealed communal chemical shift perturbation while adding the hairpin DNA and neutralization nanobody Nb2 or Nb4. Thus, these four residues were identified as hotspots to probe the activity related residues of PfuPol subsequent experiments. The CSP patterns resulting from the binding of PfuPol to Nb2, Nb4, or DNA are presented in the bottom panels of Figure 2A–C.

### Assignment of hotspots

To achieve assignments of these four hotspots, we first determine amino acid types of the identified hotspot residues. Using average chemical shifts and standard deviations from the Biological Magnetic Resonance Data Bank (BMRB) data (Ulrich *et al*, 2008) we generated residue specific chemical shift ranges on the 2D [^13^C,^1^H]-HMQC spectrum of [^13^C]-labeled PfuPol (Supplementary Figure S4). Through statistical analysis of ^1^H and ^13^C chemical shifts of identified hotspots, potential amino acid type(s) were defined using probability density distributions and Bayesian probability calculations (Figure 1B(i)), assigning the probability of the amino acid type of identified hotspot (Figure 1B(ii) and (iii)). Ali1 exhibits a 78.8% probability corresponding to Ala Cβ versus 21.2% for Met Cε (Figure 2G). Aro1 was unambiguously assigned to His Cε based on its unique chemical shift (Figure 2H). Aro2 shows 70.3% probability for Tyr Cζ, 25.3% for Trp Cδ, and 4.4% for His Cδ (Figure 2I), while aro3 has 85.1% probability for Trp Cδ and 14.9% for His Cδ (Figure 2J).

To further precisely assign these hotspots, we introduce an integrated approach with the experimental data and computational distance restraints. Firstly, we obtained the interaction patterns between different binders (DNA, Nb2, Nb4) with PfuPol from CSPs (Figure 1C(i)). Then, AlphaFold3 (Abramson *et al*, 2024) was employed to generate the structure of the complex between the experimental hairpin DNA and PfuPol, while those with nanobodies (Nb2 and Nb4) were obtained through molecular docking (Honorato *et al*, 2024). The top ten lowest energy structures of the complex of Nb2 and Nb4 with PfuPol were selected respectively. The distance (r) of each residue of PfuPol with the surface of hairpin DNA, Nb2 and Nb4 were calculated for each conformer (Figure 1C(ii) and Supplementary Figure S5). The quality of the AlphaFold3 models was validated through multiple approaches. The model for the hairpin DNA-PfuPol complexes yielded an ipTM score of 0.72, indicating good confidence, and was integrated with the crystal structure of PfuPol-DNA to derive distance information. Meanwhile, the AlphaFold3 model for each nanobody-PfuPol complexes (iPTM for PfuPol-Nb2: 0.85; iPTM for PfuPol-Nb4: 0.85) were optimized using HADDOCK2.4, with model quality assessed by the HADDOCK score (the top ten scoring structures with the lowest energy were selected). Overall, the computed ipTM scores (Supplementary Table S2) supported the validity of the AlphaFold3 models. This strengthens the integration of distance constraints with CSPs. Based on previous studies (Stark and Powers, 2008; Liu *et al*, 2017), the minimal distance between any atom of the perturbed residue and any atom of the binder conformer was assumed to be <6.0 Å. For complex models generated by AlphaFold3 and HADDOCK2.4, a positional uncertainty of 1.65 Å was applied (Abramson *et al*, 2024; Honorato *et al*, 2024). Consequently, an upper distance limit of 7.65 Å was adopted as the reference threshold for binders induced CSPs on PfuPol.

By integrating the probabilistic assessment of amino acid types for each hotspot with their spatial distances to the hairpin DNA, Nb2, and Nb4, a systematic analysis was conducted. For the ali1 hotspot, residue A684 exhibited distances of 4.8 Å, 4.6 Å, and 4.2 Å to the hairpin DNA, Nb2, and Nb4, respectively (Figure 2K and Supplementary Table S3), which aligns well with the observed CSP pattern. This clear distance profile strongly supported its assignment. In contrast, other candidate residues across all hotspots often showed contradictory trends either displaying significant CSPs despite long distances or showing no notable CSPs despite being in close proximity (Supplementary Figure S6A and Table S3). Although spatial proximity does not necessarily guarantee CSPs generation, and residues distant from binding interfaces may occasionally display CSPs due to indirect effects, such cases require stronger assumptions and more complex interpretations. The CSP data for A684 were fully explicable without resorting special hypotheses (Figure 2A–C, bottom panels). Similarly, aro1, aro2, and aro3 were evaluated using the same integrated approach, taken the possibility of each hotspot belonging to a certain amino acid type, CSP patterns, and distances of hotspots to hairpin DNA, Nb2 and Nb4, all this information together (Figure 1C(iii)). The consolidated evidence supported the of ali1 to Cβ of A684 (Figure 2K), aro1 to Cε of H680 (Figure 2L), aro2 to Cζ of Y393 (Figure 2M), and aro3 to Cδ of W624 (Figure 2N and Supplementary Figure S6L–M). Detailed assignment parameters are provided in Supplementary Figure S6 and Table S3.

### Characterization of assigned hotspots

All four assigned hotspots (Y393, W624, H680, and A684) were mapped onto the PfuPol structural model (Figure3A). Notably, Y393 residue located within the conserved “Y-GG/A” motif of a flexible loop bridging the 3’-5’ exonuclease and polymerase domains. Although not a direct catalytic residue, this motif modulates the polymerase-exonuclease activity balance (Truniger *et al*, 1996; Böhlke *et al*, 2000). W624 lies within a loop region in close proximity to DNA. A684 engaged in the DNA minor groove near the active site, while H680 occupied a peripheral position of the PfuPol-DNA interaction cluster, contributing to the local electrostatic potential.

In a further step, we introduced a saturation mutagenesis library for the position Y393, W624, H680, and A684 individually to all other 19 alternatives for the evolution of PfuPol. Mutants of PfuPol was transformed into *E. coli* BL21 strain (Figure 1D(i)). Single colonies were picked from LB agar plates (Figure 1D(ii)) and cultured in 96-deepwell plates (Figure 1D(iii)). Owing to the high thermostability of PfuPol (Lundberg *et al*, 1991), we directly utilized the recombinant *E. coli* cells as the source of PfuPol in the PCR system (Figure 1D(iv)). Thermal pre-lysis of the *E. coli* resuspension reduces degradation of oligonucleotides and dNTPs by components in the lysate. The PCR reaction with *E. coli* cell expressing PfuPol mutants was performed by using a long template for amplification (Figure 1D(v)). Surprisingly, saturation mutagenesis of these hotspot residues resulted in dramatic alterations in catalytic activity, providing strong evidence for the accuracy of the assignments and confirming the functional relevance of these hotspots to PfuPol activity. Notably, for all four assigned residues Y393, H680, and A684, we screened mutants with superior performance, mainly higher yield, than the wide type PfuPol (Figure 3D and Supplementary Figure S7A) without compromising fidelity (Supplementary Table S4). In contrast, for W624, only the W624S mutant retained activity comparable to that of the wild-type enzyme (Supplementary Figure S7A). This outcome demonstrated the identified hotspots were relevant to the activity of PfuPol (Bhattacharya *et al*, 2022; Gutierrez-Rus *et al*, 2025).

**Figure 3.**
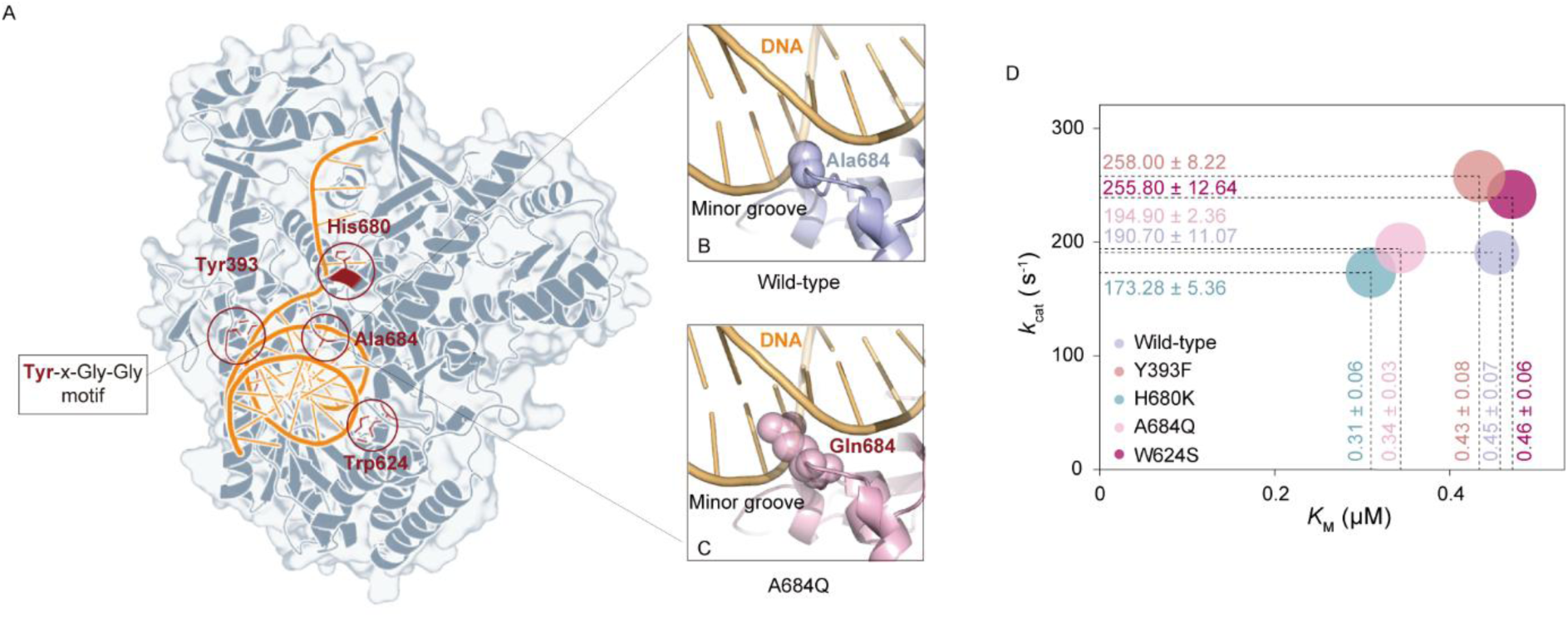
Structural and biochemical analysis of variants of PfuPol. (**A**) Structural models taken from AlphaFold3 of complex of PfuPol with a hairpin DNA. Assigned hotspots were mapped on the structure of PfuPol showing as red sticks. (**B** and **C**) Structural illustration of the DNA substrate binding pocket of the wild-type PfuPol (**B**, lightblue) and the variant PfuPol-A684Q (**C**, lightpink), predicted by AlphaFold3. The DNA substrate is shown in ball and stick representation colored by orange. The mutated residue of PfuPol and PfuPol-A684Q was drawn as spheres. All structure figures were made using PyMOL. (**D**) Comparison of kinetic parameters for PfuPol and its variants. Bubble size represents fitted k_cat_/K_M_ values. Wild-type (lightblue): PfuPol; Y393F (deepsalmon): PfuPol-Y393F; H680K (cyan): PfuPol-H680K; A684Q (pink): PfuPol-A684Q; W624S (purple): PfuPol-W624S.

To elucidate mechanisms underlying enhanced activity and further optimize PfuPol, we determined the kinetic parameters *via* Michaelis-Menten fitting (Figure1E(iii)) using an EvaGreen-based (Mao *et al*, 2007) fluorescent polymerase activity assay (Figure 1E(ii)). These parameters were subsequently analyzed in conjunction with the AlphaFold3-generated structures of PfuPol variants complexes with DNA. For PfuPol-A684Q variant, structural analysis revealed deeper penetration of the glutamine side chain into the DNA minor groove compared to wild-type (Figure 3B and C), suggesting enhanced DNA stabilization during long-template amplification and increased processivity. This result was consistent with a reduced *K*_M_ value of 0.34 ± 0.03 μM of PfuPol-A684Q variant compared with the *K*_M_ value of 0.45 ± 0.07 μM of PfuPol (Figure 3D). For PfuPol-Y393F, substitution with phenylalanine improved the activity of PfuPol in amplify the long template, whereas non-aromatic mutations abolished polymerase function (Supplementary Figure S7A), showing the essential role of aromaticity in this position. Structural comparisons with the wild-type PfuPol (Supplementary Figure S7B and C) revealed subtle conformational rearrangements and altered intramolecular interactions within the flexible loop harboring the Y-GG/A motif, consistent with the observed increased *k*_cat_ (190.7 ± 11.1 s^-1^ to 258.0 ± 8.22 s^-1^). Conversely, compared to wild-type, the PfuPol-H680K variant enhanced positive surface electrostatic potential (Supplementary Figure S7D and E), likely strengthening DNA backbone contacts and reducing *K*_M_ (0.45 ± 0.07 to 0.31 ± 0.06 μM), demonstrating dominance of charge over aromaticity at this site. In the absence of DNA, the local loop region containing W624 in the PfuPol-W624S mutant adopts a spatial conformation that is markedly different from that of the wild-type enzyme (RMSD = 3.317 Å) (Supplementary Figure S7F). Because of DNA binding, the flexible loop in the mutant becomes locked into a specific active conformation that is highly similar to the wild-type structure (RMSD = 0.556 Å) (Supplementary Figure S7G). This conformational stabilization results in a modest increase in the *k*_cat_ value of PfuPol-W624S (Supplementary Figure 3D and Table S4). Collectively, these functional enhancements validate the catalytic relevance of the hotspot assignments.

### Exploration of hotspot synergy in catalytic activity

To elucidate the underlying mechanisms of catalytic enhancement and identify optimal mutants (Figure 1E(i)), we introduced combinatorial mutagenesis of PfuPol, generating variants including three double mutants PfuPol*-*Y393F/H680K, PfuPol*-*Y393F/A684Q, and PfuPol*-*H680K/A684Q and two triple mutants PfuPol*-*Y393F/H680K/A684Q and PfuPol*-*Y393F/W624S/A684Q. We prepared these variants and determined their Michaelis-Menten kinetics of rate as a function of substrate concentration (Figure 4A). With the exception of W624S, which proved incompatible in combinatorial mutants (Supplementary Table S5), the combinatorial mutants exhibited a 2- to 10-fold modulation in catalytic efficiency relative to the wild-type enzyme. Specifically, PfuPol*-*Y393F/A684Q and PfuPol*-*Y393F/H680K/A684Q showed a comparable twofold enhancement, with catalytic efficiencies of (1.10 ± 0.25) × 10^9^ M^-1^s^-1^ and (1.03 ± 0.16) × 10^9^ M^-1^s^-1^, respectively, compared to the wild-type value of (0.52 ± 0.14) × 10^9^ M^-1^s^-1^. In contrast, the double mutants PfuPol*-*Y393F/H680K and PfuPol*-*H680K/A684Q—each containing the distal side chain mutation H680K in combination with an additional hotspot mutation—displayed a more substantial improvement of over 10-fold, achieving values of (5.81 ± 1.52) × 10^9^ M^-1^s^-1^ and (5.70 ± 1.00) × 10^9^ M^-1^s^-1^, respectively (Figure 4B), indicating strong synergistic effects. Notably, a high correlation was found between catalytic efficiency (*k*_cat_/*K*_M_) and the catalytic rate constant (*k*_cat_) across the combinatorial mutants (Figure 4C), suggesting that the mutations primarily enhance catalytic turnover rather than substrate binding. While tight substrate binding (a low *K*_M_) can confer high catalytic efficiency in principle, the reaction rate is largely dictated by *k*_cat_ under the high substrate conditions typical of many applied settings (Yu *et al*, 2022).

**Figure 4.**
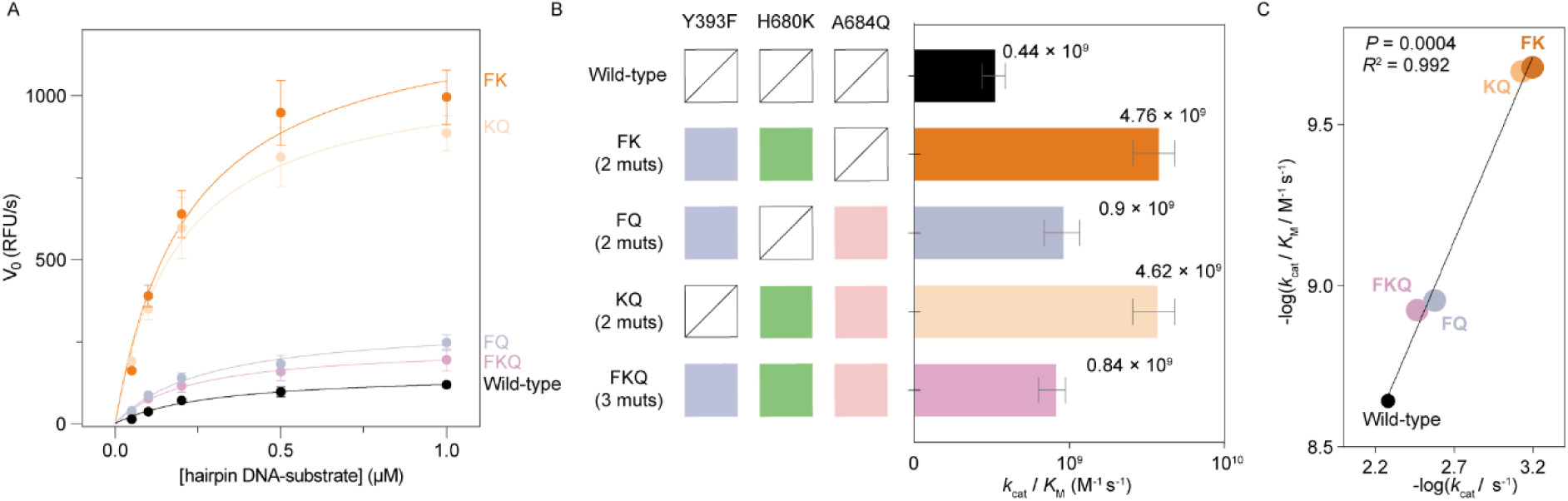
Characterization of PfuPol with combinatorial mutations at three hotspot residues. (**A**) Michaelis-Menten plots for PfuPol and its variants. (**B**) The catalytic efficiency of PfuPol and its variants. Slashes denote unmutated hotspots. Data represent the mean ± SD of 3 biological replicates. Muts, number of mutations. (**C**) Plot of logarithm of catalytic efficiency vs logarithm of catalytic rate constant including PfuPol and its variants. Wild-type (black): PfuPol; FK (brightorange): PfuPol-Y393F/H680K; FQ (lightblue): PfuPol-Y393F/A684Q; KQ (lightorange): PfuPol-H680K/A684Q; FKQ (lightpink): PfuPol-Y393F/H680K/A684Q.

## Discussion

We developed a hybrid NMR-computational approach integrating NMR experiments with computational distance constraints to map functional hotspots and guide the directed evolution of large enzymes. Applying this approach to the multi-domain PfuPol, we successfully assigned four experimentally identified hotspots, demonstrating the feasibility of integrating distance constraints with CSPs analysis for assigning functional hotspots in large enzymes. Directed evolution confirmed the catalytic relevance of these hotspots, reducing reliance on exhaustive combinatorial screening and highlighting the advantage of compact, precisely guided libraries. Critically, this approach eliminates dependency on high throughput screening. Single round rapid selection efficiently eliminates deleterious mutations while combination of the single mutant enabling cost-effective identification of optimal variants.

This study provides a streamlined pipeline for identification of hotspots in large native enzymes without complete *de novo* NMR assignments. The use of ^1^H-^13^C moieties in NMR spectroscopy extends protein analysis to mega-Dalton complexes (Mainz *et al*, 2013), enabling active site mapping in high-molecular-weight enzymes. Our approach employs distinct ^1^H-^13^C chemical shift signatures of amino acid side chains. When conventional NMR analysis becomes infeasible due to protein size, ^1^H-^13^C correlation spectra provide an effective alternative. Given potential spectral overlap challenges, distortionless enhancement by polarization transfer (DEPT) could further help to resolve methyl and methylene spectral overlap (Doddrell *et al*, 1982). Titration with substrates or neutralization nanobodies resolves functional residues *via* CSPs analysis (Tugarinov *et al*, 2007; Ruschak and Kay, 2010; Gorman *et al*, 2018). Betaine, a well-known PCR additive (Henke *et al*, 1997; Green and Sambrook, 2019) was also investigated if they acted on hotspots of PfuPol. The addition of 0.5-2.5 M Betaine (N,N,N-trimethylglycine) into the PfuPol reaction system was shown to enhance its performance in amplifying different templates with varied length (Supplementary Figure S8C–E) respectively. Meanwhile, significant CSPs of PfuPol were observed on the hotspot residues aro1, aro2, and aro3 upon the addition of 1 M betaine (Supplementary Figure S8A and B) were detected. This result showed if neutralization antibody is not available, co-factors of the target enzyme could also be used to identify the functional hotspots. Additionally, natural abundance [^13^C,^1^H]-HMQC spectra of PfuPol showed a high resolution and signal to noise ratio signals, enabling hotspots detection in unlabeled enzymes (Supplementary Figure S9), demonstrating the potential of broad application of our proposed approach.

Distance constraints derived from enzyme-binder complexes correlate with CSPs to strongly assign hotspot residues, drastically narrowing mutational search space (Bhattacharya *et al*, 2022; Gutierrez-Rus *et al*, 2025; Marshall *et al*, 2023; Wu *et al*, 2022; Xu *et al*, 2019). In structural bioinformatics, CSPs are widely recognized as a valuable source of medium to low resolution restraint information (Yu *et al*, 2017; Aguirre *et al*, 2014; Wagner *et al*, 1983). Unlike the precise geometric data provided by nuclear Overhauser effects (NOEs), CSPs offer ambiguous, qualitative, and probabilistic spatial constraints. However, they play an indispensable role in identifying the overall binding mode of protein-ligand complexes. In this work, CSP-derived restraints were used as a coarse-grained filtering step an important advancement when full resonance assignments were unavailable. Although “false negatives” (i.e., proximal residues showing no significant CSPs) can occur, clusters of residues with substantial CSPs are statistically enriched at true binding sites across the protein interface. While data from a single binder may occasionally be misleading due to random effects, the consistent CSP patterns observed for three distinct binders provide multidimensional cross-validation, making the possibility of random matches exceedingly low. This greatly strengthens the reliability of the resulting conclusions. For residues that influence catalysis through long-range effects, they might be overlooked, as distance-based filtering cannot fully exclude false-negative positions. Future integration with dynamic NMR methods or artificial intelligence aided dynamic predictions may further improve accuracy and broaden applicability to enzymes governed by allosteric regulation.

Notably, our assigned hotspots encompass the tyrosine residue within the previously reported “Y-GG/A” motif of Family B polymerases (Truniger *et al*, 1996; Böhlke *et al*, 2000), which aligns with our Y393 hotspot, demonstrating the method’s reliability in identifying residues that indirectly regulate active sites. While A684 directly engages DNA and Y393 modulates activity allosterically, H680 occupies a peripheral interaction cluster. These results demonstrate the capacity of our method to probe residues proximal to functional interfaces. Variants characterization revealed some minor functional trade-offs: Lysine substitution at PfuPol-H680K improved pH stability over wild-type histidine, but arginine substitution abolished activity, indicating electrostatic interactions with DNA must remain balanced within a defined range (Supplementary Figure S7A). Hotspot-guided evolution generated variants with elevated *k*_cat_ values, diverging from traditional directed evolution strategies that typically enhance catalytic efficiency by reducing *K*_M_ (Bhattacharya *et al*, 2022). PCR assays showed the PfuPol-Y393F variant enhanced amplification efficiency across diverse template lengths (Supplementary Figure S10A), PfuPol-H680K improved specificity for select templates (Supplementary Figure S10C), and PfuPol-A684Q boosted amplification of high-GC templates (Supplementary Figure S10B). Further enhancement of the activity of PfuPol *via* combinatorial mutagenesis also confirmed the non-redundant roles of identified hotspots, demonstrating the high efficiency of our proposed approach. The W624S mutation does not completely eliminate enzymatic activity, but it increases the flexibility and induces conformational changing in the corresponding loop region under apo conditions (Supplementary Figure S7F and G). This enhanced flexibility may allow the loop to sample multiple conformational states, making it more challenging to maintain the proper architecture in the absence of substrate. DNA binding appears to compensate for the loss in intrinsic stability caused by the mutation. Thus, W624 may act as a critical “switching” residue that helps maintain the local conformation of PfuPol in the substrate free state, while the functional catalytic conformation is largely induced and stabilized by substrate binding.

In summary, this hybrid NMR-computational framework markedly broadens the utility of NMR spectroscopy in guiding the directed evolution of large enzymes. By integrating NMR methods with computational analysis, it eliminates the need for full *de novo* NMR assignments, thereby advancing enzyme engineering toward greater catalytic efficiency and functional diversity.

## Material and methods

### Cloning, expression, and purification of proteins

Genes of PfuPol was codon optimized and synthesized by Sangon Biotech Co., Ltd. (China) and cloned in a modified pET16b vector (Novagen). The mutants of PfuPol were directly obtained through library screening (see Library construction) or generated via site directed mutagenesis PCR (Supplementary Table S1). The genes of nanobodies Nb2 and Nb4 were cloned in a modified pET22b vector containing N-terminal His-tag (Tao *et al*, 2024). The respective plasmid was transformed into BL21 (DE3) competent cells (Novagen), grown in Luria Bertani (LB) broth or M9 medium supplemented with 100 μg/mL ampicillin and induced for expression with a final concentration of 0.5 mM isopropyl-D-1-thiogalactopyranoside (IPTG) at an optical density (600 nm) of 0.6 for 5 hours at 37 °C. Uniformly [^13^C]-labeled protein was prepared by growing cells in M9 medium containing [U-^13^C] glucose (3 g/liter) (He *et al*, 2015). Cells were harvested by centrifugation at 5000g for 25 min at 4 °C.

For the purification of polymerase, the pellet was resuspended in 50 mL of Lysis Buffer A (50 mM Tris-HCl pH 9.0, 300 mM KCl, 0.1 mM PMSF and 0.01 mg/mL DNase I). Afterwards, cells were lysed by a high-pressure homogenizer and the pellet was separated by centrifugation at 27,000×g for 30 min at 4 °C. The collected supernatant was incubated for 30 minutes at 70 °C to precipitate cellular proteins of *E. coli*, followed by one more round of centrifugation at 27,000×g for 25 min. The supernatant was then diluted 6-fold and loaded onto a 5 mL HiTrap Heparin column (GE Bioscience) that was pre-equilibrated with Heparin Buffer (50 mM Tris-HCl pH 8.0, 50 mM KCl). The polymerase was eluted with a 0–500 mM KCl gradient (50 mM Tris-HCl pH 8.0). The eluted protein was immediately buffer exchanged via a PD-10 column (GE Bioscience) to the Storage Buffer (25mM Tris-HCl pH 7.4, 50 mM KCl, 0.1 mM EDTA, 1 mM DTT, 50% glycerol) and stored at –20 °C.

For the purification of nanobody, the cell pellet was resuspended in 50 mL of Lysis Buffer B (20 mM Tris-HCl pH 7.4, 150 mM NaCl, 0.1 mM DTT, 0.1 mM PMSF and 0.01 mg/mL DNase I). Afterwards, cells were lysed by a high-pressure homogenizer and the pellet was separated by centrifugation at 27,000×g for 30 min at 4 °C. The supernatant containing each target protein was loaded onto a Ni-NTA (nitrilotriacetic acid) column (Qiagen) which was pre-equilibrated with Lysis Buffer B. The column was washed with 100 mL of Wash Buffer (Lysis Buffer B with 30 mM imidazole), and eluted with 25 mL Elution Buffer C (20 mM Tris-HCl pH 7.4, 150 mM NaCl, 0.1 mM DTT, 300 mM imidazole). The eluted fraction was then fractionated on a size exclusion Superdex-75 column (GE Bioscience) equilibrated in SEC Buffer (20 mM Tris-HCl pH 7.4, 150 mM NaCl, 0.1 mM DTT, 0.1 mM EDTA) to further purify the proteins. The proteins (Nb2 and Nb4) were concentrated and store at –80 °C for further usage.

### NMR spectroscopy

All NMR experiments of the PfuPol were performed in the NMR buffer (20 mM NaH_2_PO_4_/Na_2_HPO_4_ pH 7.4, 35 mM NaCl, 0.02% NaN_3_, 8% D_2_O). If not specified, all spectra were collected at 298K on Bruker Avance 600 MHz NMR spectrometer equipped with the cryogenically cooled probe. 2D [^13^C,^1^H]-HMQC spectra of 50 μM [^13^C]-labeled PfuPol were recorded in the absence and presence of 50 μM hairpin DNA or 75 μM Nb2 or 75 μM Nb4 to monitor the chemical shift change for identifying the hotspots. For all NMR spectra, 1024 complex points were recorded in the direct dimension and 154 complex points were recorded in the indirect dimension. Shifted sine bell and Polynomial baseline correction were applied in all dimensions. NMR data were processed and analyzed in CcpNmr Analysis (Vranken *et al*, 2005). The combined chemical shift perturbation (Δδ_comb_) was calculated for each residue using the formula:

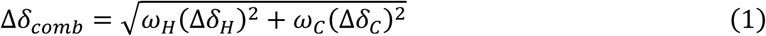

where, *Δδ*_H_ and *Δδ*_C_ are chemical shift changes (in ppm) in ^1^H and ^13^C dimensions, respectively, and *ω*_H_ and *ω*_C_ are normalization factors (*ω*_H_ = 1.00, *ω*_C_ = 0.34 for aliphatic or 0.07 for aromatic).

### Assignments of hotspots

The peaks from the 2D [^13^C,^1^H]-HMQC spectra of PfuPol, showing common chemical shift perturbation upon the titration of hairpin DNA, nanobody Nb2 and Nb4 individually were identified as the hotspots. To assign the hotspots, we first compared the chemical shifts of hotspots with the statistics values of amino acids from the Biological Magnetic Resonance Bank (BMRB) (Ulrich *et al*, 2008) database to calculate the possibility of these hotspots belonging to a certain amino acid type. The probability density function *f*(a, b) for a hotspot residue belongs to a certain amino acid types is given by:

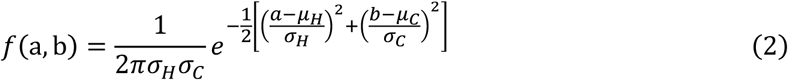

where a is the chemical shift of the hotspot residue in the Hydrogen dimension, b is the chemical shift of the hotspot residue in the carbon dimension, *μ*_*H*_ is the mean chemical shift in the Hydrogen dimension for the amino acid type, *σ*_*H*_ is the standard deviation of the chemical shift in the Hydrogen dimension for the amino acid type, *μ*_*C*_ is the mean chemical shift in the carbon dimension for the amino acid type, *σ*_*C*_ is the standard deviation of the chemical shift in the carbon dimension for the amino acid type. Using Bayes’ theorem, the probabilities that the hotspot belongs to certain amino acid type are:

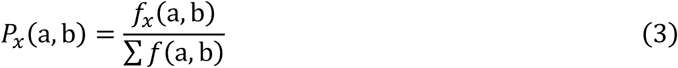

where *f*_*x*_(a, b) is the probability density value for the hotspot residue belonging to amino acid type *x*, ∑ *f*(a, b) is the total probability density values for all possible amino acid types of the hotspot residue.

With the possibility of the hotspot belonging to which amino acid, we then applied the integrated methods with AlphaFold3 and HADDOCK2.4 to assign the hotspot. The structure of PfuPol-hairpin DNA complex was generated by AlphaFold3 (Abramson *et al*, 2024) Based on residue level confidence scores (pLDDT) and interface pLDDT (I-pLDDT) scores, the top 5 models were selected for later usage. The neutralization nanobodies Nb2 and Nb4 were docked with PfuPol *via* HADDOCK with a previously described protocol (Honorato *et al*, 2024) 100 docking models for each PfuPol-nanobody complex were generated. According to haddock-score, the ensemble of 10 conformations with the lowest energy were selected. Interatomic distances (r) between residues of PfuPol and binding partners, specifically DNA or nanobodies Nb2 and Nb4, were measured using PyMOL. The minimum distance between any atom of the perturbed residue and any atom of the binder conformer was set to less than 6.0 Å, following previous studies (Stark and Powers, 2008; Liu *et al*, 2017). For complex structures generated by AlphaFold3 and HADDOCK2.4, a positional uncertainty of 1.65 Å was incorporated to account for model variability (Abramson *et al*, 2024; Honorato *et al*, 2024). Thus, an upper distance threshold of 7.65 Å was defined as the reference criterion for identifying binder-induced CSPs in PfuPol. The distance (r) between each PfuPol residue of the assigned amino acid type with the corresponding binder (hairpin DNA, Nb2, or Nb4) was evaluated against the upper distance threshold of 7.65 Å. Residues that consistently had a distance with hairpin DNA, Nb2, and Nb4 less than 7.65 Å were assigned to the identified hotspot.

### Library construction

Saturation mutagenesis library of PfuPol was created *via* site-directed PCR with wild-type PfuPol plasmid cloned in pET16b vector as the template. Degenerate primer pairs (containing an NNK codon at the position to be mutated) were designed to mutate residues at positions 393, 680 and 684. Primer sequences are provided in Supplementary Table S1. Site-directed mutagenesis was done with FasHifi Super DNA polymerase (Swiss Affinibody Lifescience Co., Ltd.) using standard protocol. After the PCR step, the template was digested with Dpn I restriction enzyme and then inactivated at 80 °C, followed by transformation into *E. coli* BL21 (DE3) competent cells.

### Library Screening

*E. coli* BL21 (DE3) cells containing PfuPol mutants were plated on LB agar with 100 mg/mL ampicillin. A single colony was picked and inoculated in a 96 deep-well plate (with 2 mL LB medium and 100 mg/mL ampicillin per well), sealed using a breath-easy membrane. Cultures were incubated at 37 °C induced for expression with a final concentration of 0.3 mM IPTG at an optical density (600 nm) of 0.8 for 3 hours at 220 rpm. Cells were harvested by centrifugation at 3500g for 20 min at 4 °C. Pellets were resuspended in ddH_2_O, centrifuged again, and the supernatant was discarded. Cells were resuspended in ddH_2_O, and their OD_600_ values were measured. Subsequent stepwise dilution into ddH_2_O enabled determination of the optimal *E. coli* cells concentration for PCR reactions using wild-type PfuPol. We established that adding 1 μL of *E. coli* cells suspension (OD_600_ = 0.33) to 20 μL reaction volumes yielded maximal activity. Screening was performed in two stages: first, short-fragment amplification (LucB primer sequences in Supplementary Table S1) promptly eliminated clones lacking detectable activity; second, long-fragment pressure screening (Long primer sequences in Supplementary Table S1) identified functionally robust variants. Specifically, first, incubation at 98 °C for 6 min to lyse cells for releasing the highly thermostability of PfuPol proteins. Then, followed by 34 thermal cycles of 98 °C for 15 s, 60 °C for 15 s, and 72 °C for 20 sec to 5 min depending on the target DNA length (1 kb/min). The reaction buffer contained 50 mM Tris-HCl pH 8.8, 10 mM KCl, 2.5 mM MgCl_2_, 0.2 mM dNTPs, 1 M betaine, 0.001% BSA and 0.05% Triton X-100. Afterwards, 4 μL of the PCR product was mixed with the loading dye and applied onto a 1% agarose-TAE gel containing 1× GenRed nucleic acid gel stain (Genview, China). After electrophoresis, the gel was photographed under ultraviolet light. The gels were analyzed by ImageJ to qualify the intensity of PCR product bands. The band intensities of the wild-type were used for normalization, and the relative activities of the variants were calculated. The heatmap of relative activities was generated using the software GraphPad Prism 8.0 (GraphPad Software, Inc.). Plasmids extracted from the colonies demonstrating improved activity were sequenced by Sanger sequencing (Sangon Biotech Co., Ltd.) to determine the identities of beneficial mutations.

### Steady-state kinetic analyses

Steady-state kinetic data were collected and analyzed using the EvaGreen-based fluorometric polymerase activity assay (Mao *et al*, 2007). The hairpin DNA (T) (sequence in Supplementary Table S1) was prepared in advance by heating at 98 °C for 5 min and annealing on the ice for 30 min. Different amounts of hairpin DNA (T) at the indicated concentrations were then used as the template to mixed with dNTPs, 8 nM polymerase respectively. MgCl_2_ was added to initiate the DNA synthesis at 72 °C. For PfuPol and its variants, the reaction buffer contained 25 mM Tris-HCl pH 8.8, 10 mM KCl, 2.5 mM MgCl_2_, 1× EvaGreen dye, 0.2 mM dNTPs, and 0.05% Triton X-100. The fluorescence curve was plotted against the time, where the value of the fluorescence was derived by subtracting the background fluorescence value of each reaction. The initial rate of each reaction with different amounts of template was obtained by calculating the first derivative of the fluorescence curve. The kinetics parameters of *k*_cat_ and *K*_M_ were generated by fitting the Michaelis-Menten kinetic equation with the software GraphPad Prism 8.0. The reported values are the average of a biological duplicate each with three technical replicates.

