## Supplementary Information for "Functional hotspots identification *via* a hybrid NMR-computational approach facilitating directed evolution of large enzyme"

**This PDF file includes:**

Supplementary Methods
Supplementary Figure S1 to S10
Supplementary Table S1 to S5

### Supplementary Methods

**DNA polymerase assay.** The hairpin DNA (T) was prepared in advance by heating at 98 °C for 5 min and annealing on the ice for 30 min. 20 μl reactions were prepared by mixing buffer (10 mM Tris-HCl pH 8.8, 10 mM KCl, 2.5 mM $MgCl_2$, 200 μM dNTPs, and 0.1% Triton X-100) with 1 μM hairpin DNA (T) and 10 nM PfuPol in the presence and absence of 10 nM Nb2 or Nb4 respectively. $MgCl_2$ was added to initiate the DNA synthesis at 72°C. Following incubation at 72 °C for 1 h, reactions were quenched on ice by addition of 20 μL of 2× denaturing loading buffer (95% deionized formamide, 0.05% SDS, 20 mM EDTA 0.025% (w/v) bromophenol blue, 0.025% (w/v) xylene cyanol FF), before heating at 98 °C for 5 min. Products were resolved on 15% (w/v) Urea-PAGE gel. after electrophoresis, the gel was stained by SYBR Gold and imaged under ultraviolet light.

**3′→5′ Exonuclease assay.** Pretreatment of hairpin DNA (E) (Table S1) is as described above. 20 μl reactions were prepared by mixing buffer (10 mM Tris-HCl pH 8.8, 10 mM KCl, 2.5 mM $MgCl_2$, and 0.1% Triton X-100) with 1 μM hairpin DNA (E) and 10 nM PfuPol in the presence and absence of 10 nM Nb2 or Nb4 or 200 μM dNTPs respectively. Following incubation at 50 °C for 1 h, reactions were quenched on ice by addition of 20 μL of 2× denaturing loading buffer, before heating at 98 °C for 5 min. Products were resolved on 15% (w/v) Urea-PAGE gel. After electrophoresis, the gel was stained by SYBR Gold and imaged under ultraviolet light.

**Fluorescence based thermal shift assay.** The thermostability experiments were measured via QuantStudio 3 Real-Time PCR instrument from Thermo Fisher Scientific. The commercial dye SYPRO Orange (Sigma) was used to monitor the thermodynamic stability of both proteins. The melting temperature was programmed from 25 °C to 98 °C with a 0.15 °C temperature increment. Each reaction was performed in a final volume of 20 μL containing 50 mM Tris-HCl pH 7.4, 10 mM KCl, 10× SYPRO Orange and 10 μM of every individual protein. The signal was collected per second. The first derivative of the fluorescence signal curve was calculated using GraphPad Prism 8.0 (GraphPad Software, Inc.).

**Endpoint PCR.** 20 μL reactions were prepared by mixing buffer (50 mM Tris-HCl pH 8.8, 10 mM KCl, 6 mM ammonium sulfate, 2 mM $MgCl_2$, 0.05% Triton X-100, 0.001% BSA) with 25 nM PfuPol or its variants, 4 ng (plasmid DNA) or 40 ng (cotton gDNA) or 45 ng (rice cDNA) templates, 0.5 μM of each primer (*SI Appendix*, Table S1), 0.2 mM dNTP and varying amounts of additives (see figure legends). Amplification reactions were performed using a thermos cycler (C1000 Touch, Bio-rad): First, incubation at 95 °C for 3 min; followed by 33 thermal cycles of 95 °C for 30 s, 55 °C for 30 s, and 72 °C for 45 sec to 10 min depending on the target DNA length (1 kb/min); Afterwards, 4 μL of the PCR product was mixed with the loading dye and applied onto a 1% agarose-TAE gel containing 1× GenRed nucleic acid gel stain (Genview, China). After electrophoresis, the gel was photographed under ultraviolet light. The gels were analyzed by ImageJ (National Institutes of Health, NIH) to qualify the intensity of PCR product bands. Plots of the gel band intensity were generated with the software GraphPad Prism 8.0.

**PCR Fidelity assay.** The circular pUC19 plasmid was linearized by digestion with EcoR I. The linearized pUC19 plasmid were amplified by the PfuPol or its variants in the buffer containing 25 mM Tris-HCl pH 8.8, 10 mM KCl, 6 mM $(NH_4)_2SO_4$, 2 mM $MgCl_2$, 200 μM dNTPs, and 0.05% Triton X-100. The concentrations of the DNA polymerase and the linearized plasmid were 50 nM and 2 nM, respectively. Amplification reactions were performed using a thermos cycler (C1000 Touch, Bio-rad): First, incubation at 95 °C for 2 min; followed by 26 thermal cycles of 95 °C for 15 s, 55 °C for 15 s, and 72°C for 3 min. The linearized pUC19 plasmid PCR product was examined using 1% agarose gel electrophoresis and recovered by gel excision, following by a ligation assay with the T4 DNA ligase. Afterwards, 2.5 μL of the ligation product was transformed into *Turbo* competent cells (NEB). Transformed cells were plated on LB agar with 100 mg/mL ampicillin, 1 mg/mL X-Gal and 1.5 mM IPTG. Blue/white colonies were counted to calculate the error rates using the equation: $ER = mf/(bp \times d)$, Where, $mf$ is the mutation frequency determined by dividing the total number of white plaques by the total number of plaques. $d$ is the number of DNA duplications and $b$ is the effective target size of the linearized pUC19 plasmid, which is 323 bp.

**NMR spectroscopy for unlabeled enzymes.** NMR experiments of unlabeled PfuPol were performed in the buffer containing 20 mM $NaH_2PO_4/Na_2HPO_4$ pH 7.4, 35 mM NaCl, 0.02% $NaN_3$, 8% $D_2O$. 2D [$^{13}C$,$^{1}H$]-HMQC spectra of 1 mM unlabeled PfuPol were recorded at 298K on Bruker Avance 800 MHz NMR spectrometer equipped with the cryogenically cooled probe. For all NMR spectra, 1024 complex points were recorded in the direct dimension and 154 complex points were recorded in the indirect dimension.

### Supplementary Figures

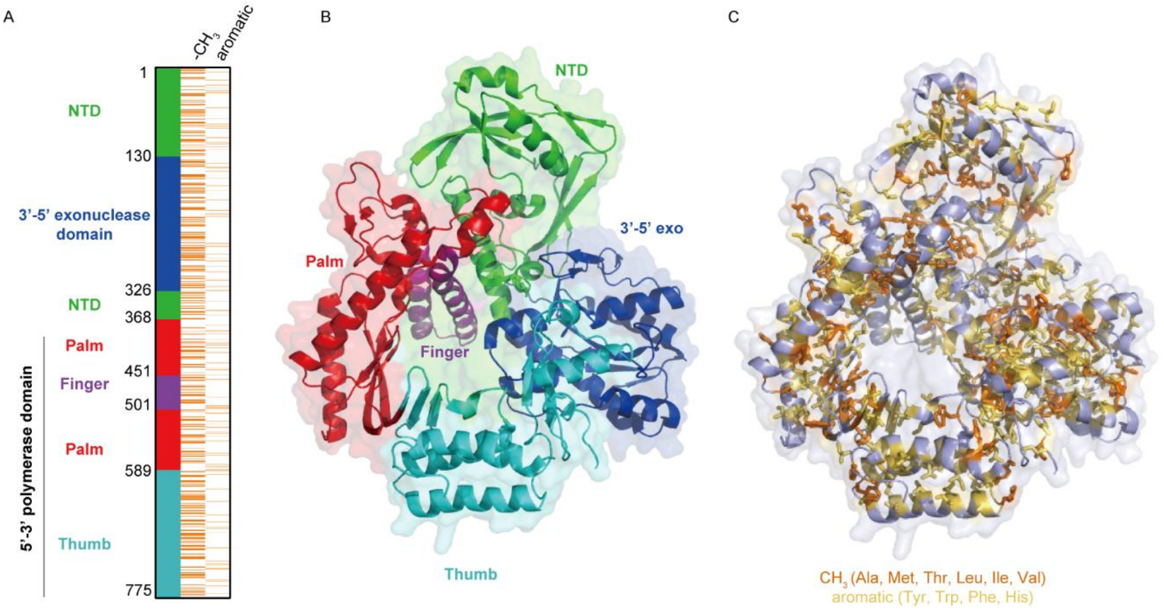


**Figure S1.** (**A**) Heatmap of the domain architecture of PfuPol and the distribution of methylated residues and aromatic residues. (**B**) The structure model is composed of domains and subdomains, which are N-terminal domain (NTD, green), 3'-5' exonuclease domain (3'-5' exo, dark blue), 5'-3' polymerase domain including the Palm (red) and Fingers (violet) subdomains and the Thumb domain (sky blue). (**C**) Residues with methyl groups and aromatic residues are represented in the structure model by orange and golden yellow, respectively.

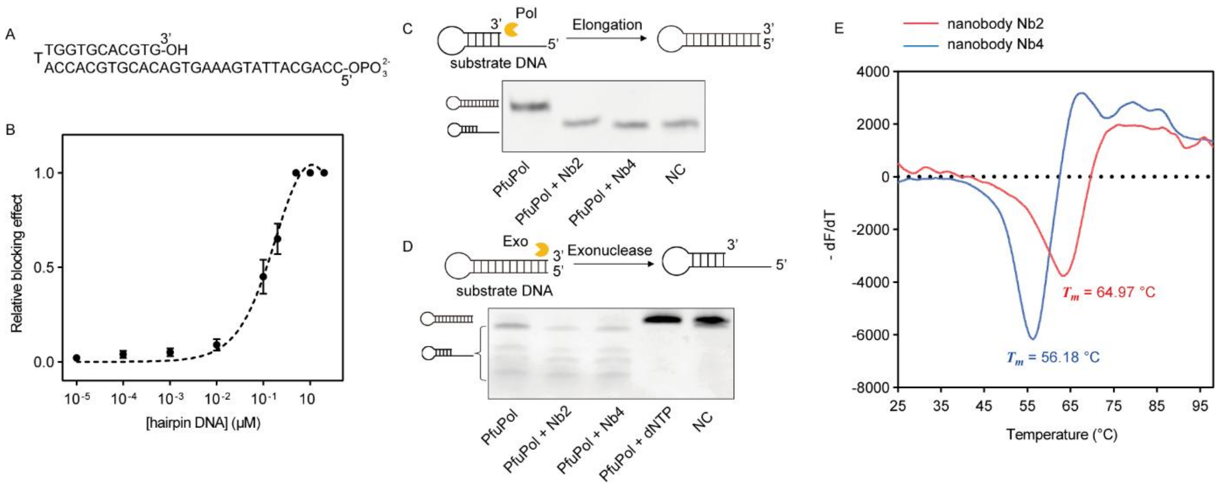


**Figure S2.** (**A**) Hairpin DNA (T) sequence derived from a previously reported variant. (**B**) The hairpin DNA (T) competes for blocking DNA polymerase activity of PfuPol (**C**) Nanobody Nb2 and Nb4 block the DNA polymerase activity of PfuPol. The lanes of PCR products with either nanobody Nb2 or Nb4 showed polymerase blocking activity, where no elongation of hairpin DNA occurs. (**D**) Nanobody Nb2 and Nb4 cannot block the exonuclease activity of PfuPol. The lanes containing PCR products with either nanobody Nb2 or Nb4 showed no detectable exonuclease blocking activity. (**E**) Thermostability of nanobody Nb2 (red) and Nb4 (blue). NC: negative control.

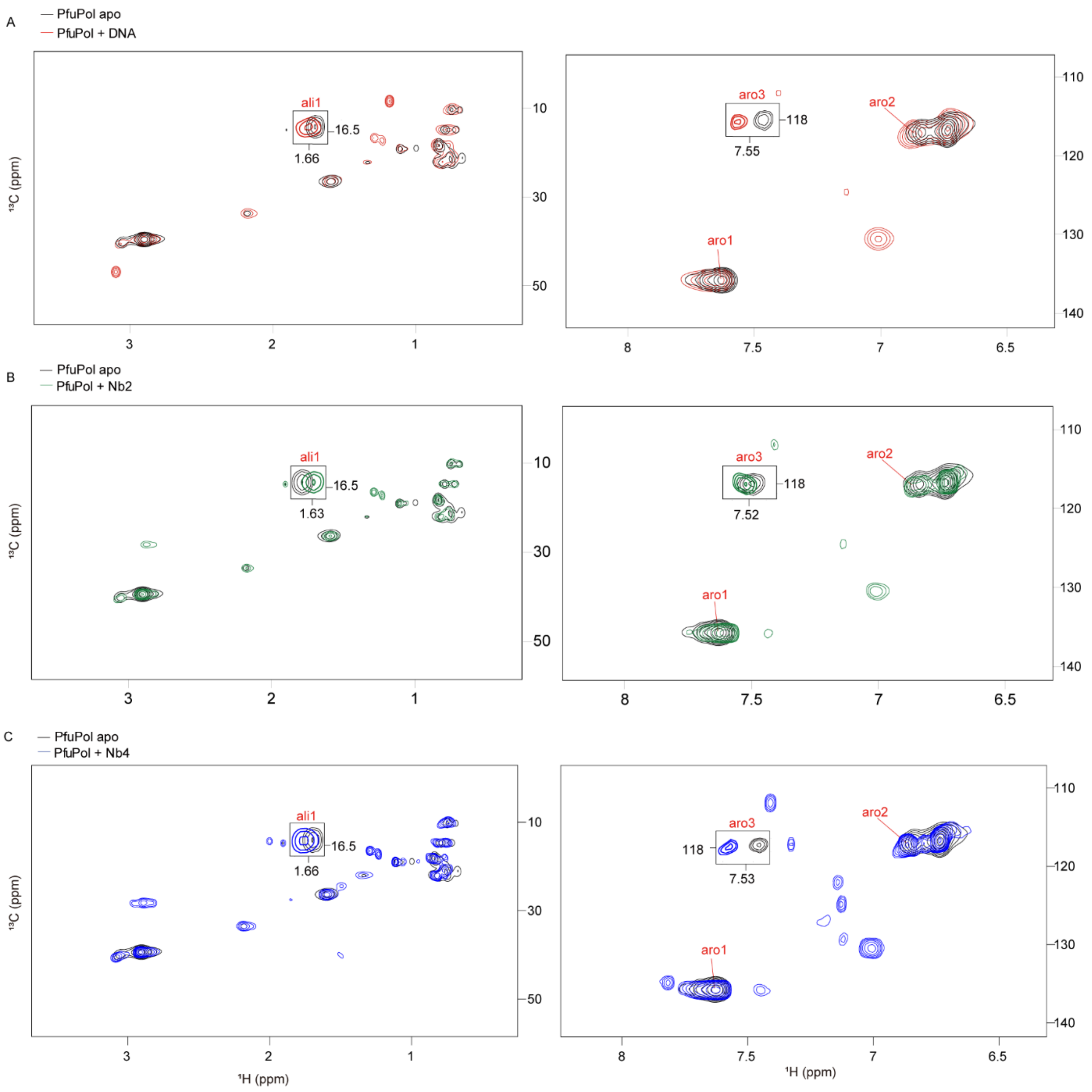


**Figure S3.** 2D [$^{13}$C,$^{1}$H]-HMQC spectra of 50 µM [$^{13}$C]-labeled PfuPol acquired on the 600 MHz ($^{1}$H frequency) spectrometer at 298 K in the absence and presence of 50 µM hairpin DNA (T) (**A**), or 75 µM nanobody Nb2 (**B**) or 75 µM nanobody Nb4 (**C**). The spectrum was processed with a positive base level of 25000.

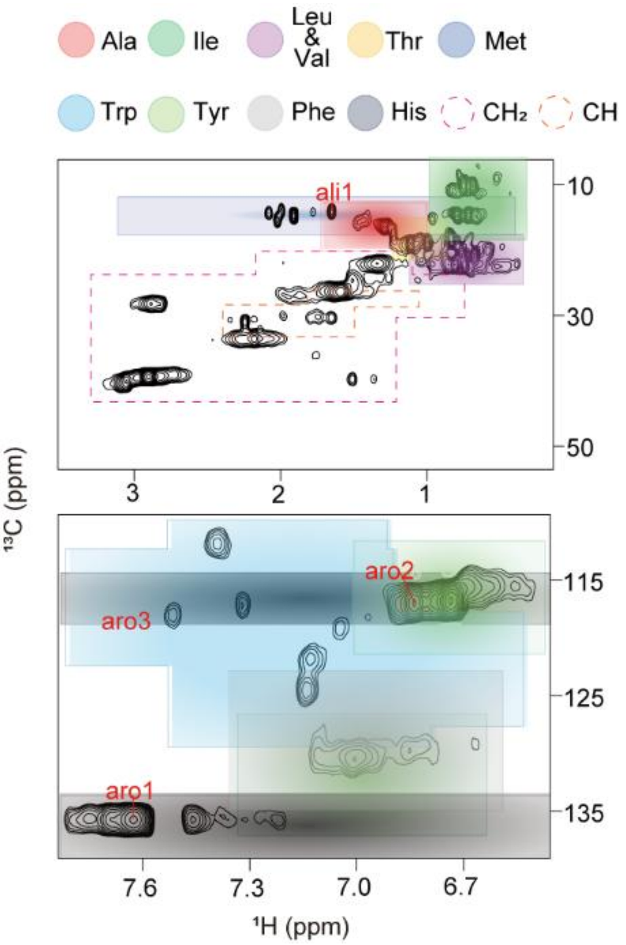


**Figure S4.** Two-dimensional [$^{13}$C,$^{1}$H]-HMQC spectra of 50 μM [$^{13}$C]-labeled PfuPol were acquired on the 600 MHz spectrometer at 298 K, where identified hotspots are labeled in red (ali1, aro1, aro2, and aro3). The spectrum was processed with a positive base level of 2500 to enhance the visualization of all hotspots. Two-dimensional [$^{13}$C,$^{1}$H]-HMQC spectra of 50 μM [$^{13}$C]-labeled PfuPol acquired on the 600 MHz spectrometer at 298 K, where the resonances of the methyl groups and aromatic groups are colored by different colors. The chemical shift range of each amino acid is represented by a rectangle centered at the mean $^{1}$H and $^{13}$C chemical shift values. The width and height of each rectangle correspond to twice the standard deviation of the $^{1}$H and $^{13}$C chemical shifts, respectively, where mean values and standard deviation of $^{1}$H and $^{13}$C are from the Biological Magnetic Resonance Bank. The terminal methyl groups of aliphatic amino acid side chains exhibit relatively dispersed carbon chemical shifts, except for Val and Leu. Based on the $^{1}$H and $^{13}$C chemical shift, the peaks of aliphatic methyl groups can be assigned to no more than three amino acid types, except threonine. In contrast, aromatic amino acids have a broader $^{13}$C chemical shift distribution, significantly reducing peak overlap. Thus, the peaks of aromatic groups can be assigned to no more than two amino acid types, except histidine. Ala, alanine-$CH_3$ (red); Ile, isoleucine-$CH_3$ (pale green); Leu & Val, leucine-$CH_3$ & valine-$CH_3$, (pink); Thr, threonine-$CH_3$ (yellow); Met, methionine-$CH_3$ (blue); Trp, tryptophan-Ind (pale blue); Tyr, tyrosine-PhO (green); Phe, Phenylalanine-Ph (grey); His, histidine-Im (light black); $CH_2$, methylene group of residues (pink dashed line); CH, methine group of leucine and valine (orange dashed line). $CH_3$, methyl; Ind, indole; PhO, phenyl-OH; Ph, phenyl; Im, Imidazole.

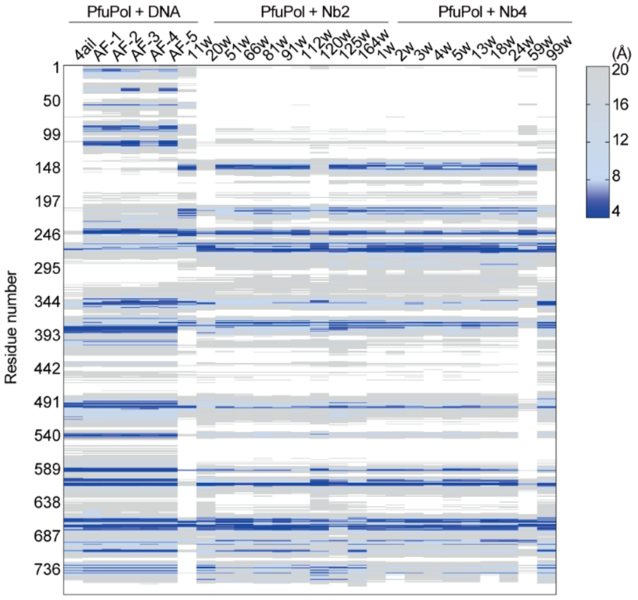


**Figure S5.** Hierarchical clustering of interaction distances of PfuPol complexes. Sequence coverage plot (heatmap) visualizes the intermolecular distance between PfuPol residues (y-axis) and bound ligands (DNA, Nb2, or Nb4) in various in silico models. For the PfuPol-DNA complex, the x-axis includes the experimental crystal structure (4ail) and five predicted models (AF-1-5) by AlphaFold3. Across all conditions, each column represents a representative structure from a cluster of conformations, with labels serving as arbitrary cluster identifiers. Interaction intensity is represented by a blue color gradient (right scale), where darker shades of blue indicate higher values (~20 Å), and lighter shades indicate lower values (~4 Å). Residues at a distance of 30 Å or greater are indicated in white. This analysis identifies the binding interface residues of PfuPol with different substrates.

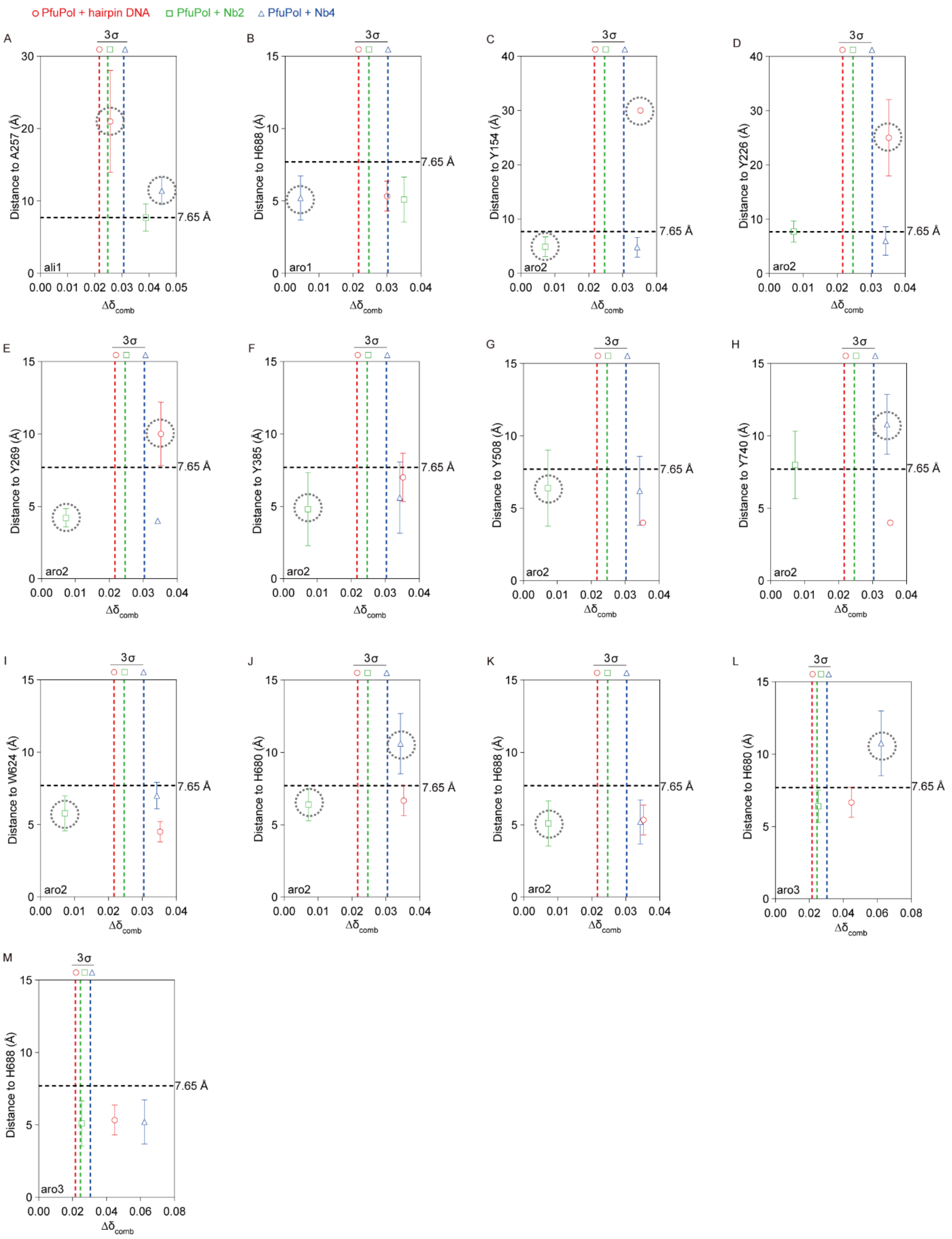


**Figure S6.** The plot of the minimum distance (r) between residues in PfuPol and their interaction partners (DNA, Nb2, Nb4) versus the chemical shift perturbation ($\Delta\delta_{comb}$) induced by the hotspots and the interaction partners (DNA, Nb2, Nb4). (**A**) Residue A257 was excluded as a candidate for ali1, as it shows significant $\Delta\delta_{comb}$ but excessive distances (> 7.65 Å) to DNA and Nb4. (**B**) Residue H688 was excluded as a candidate for aro1, as it shows insignificant $\Delta\delta_{comb}$ but exhibits spatial proximity (minimum distance < 7.65 Å) with Nb4. (**C**) Residue Y154 was excluded as a candidate for aro2, as it shows insignificant $\Delta\delta_{comb}$ but exhibits spatial proximity (minimum distance < 7.65 Å) with Nb2, and it shows significant $\Delta\delta_{comb}$ but excessive distances (> 7.65 Å) to DNA. (**D**) Residue Y226 was excluded as a candidate for aro2, as it shows significant $\Delta\delta_{comb}$ but excessive distances (> 7.65 Å) to DNA. (**E**) Residue Y269 was excluded as a candidate for aro2, as it shows insignificant $\Delta\delta_{comb}$ but exhibits spatial proximity (minimum distance < 7.65 Å) with Nb2, and it shows significant $\Delta\delta_{comb}$ but excessive distances (> 7.65 Å) to DNA. (**F**) Residue Y385 was excluded as a candidate for aro2, as it shows insignificant $\Delta\delta_{comb}$ but exhibits spatial proximity (minimum distance < 7.65 Å) with Nb2. (**G**) Residue Y508 was excluded as a candidate for aro2, as it shows insignificant $\Delta\delta_{comb}$ but exhibits spatial proximity (minimum distance < 7.65 Å) with Nb2. (**H**) Residue Y740 was excluded as a candidate for aro2, as it shows significant $\Delta\delta_{comb}$ but excessive distances (> 7.65 Å) to Nb4. (**I**) Residue W624 was excluded as a candidate for aro2, as it shows insignificant $\Delta\delta_{comb}$ but exhibits spatial proximity (minimum distance < 7.65 Å) with Nb2. (**J**) Residue H680 was excluded as a candidate for aro2, as it shows insignificant $\Delta\delta_{comb}$ but exhibits spatial proximity (minimum distance < 7.65 Å) with Nb2, and it shows significant $\Delta\delta_{comb}$ but excessive distances (> 7.65 Å) to Nb4. (**K**) Residue H688 was excluded as a candidate for aro2, as it shows insignificant $\Delta\delta_{comb}$ but exhibits spatial proximity (minimum distance < 7.65 Å) with Nb2. (**L**) Residue H680 was excluded as a candidate for aro3, as it shows significant $\Delta\delta_{comb}$ but excessive distances (> 7.65 Å) to Nb4. (**M**) The assignment of aro3 to the residue H688 had minimum distance with all three binders (DNA, Nb2, Nb4) less than 7.65 Å, meanwhile showing significant $\Delta\delta_{comb}$.

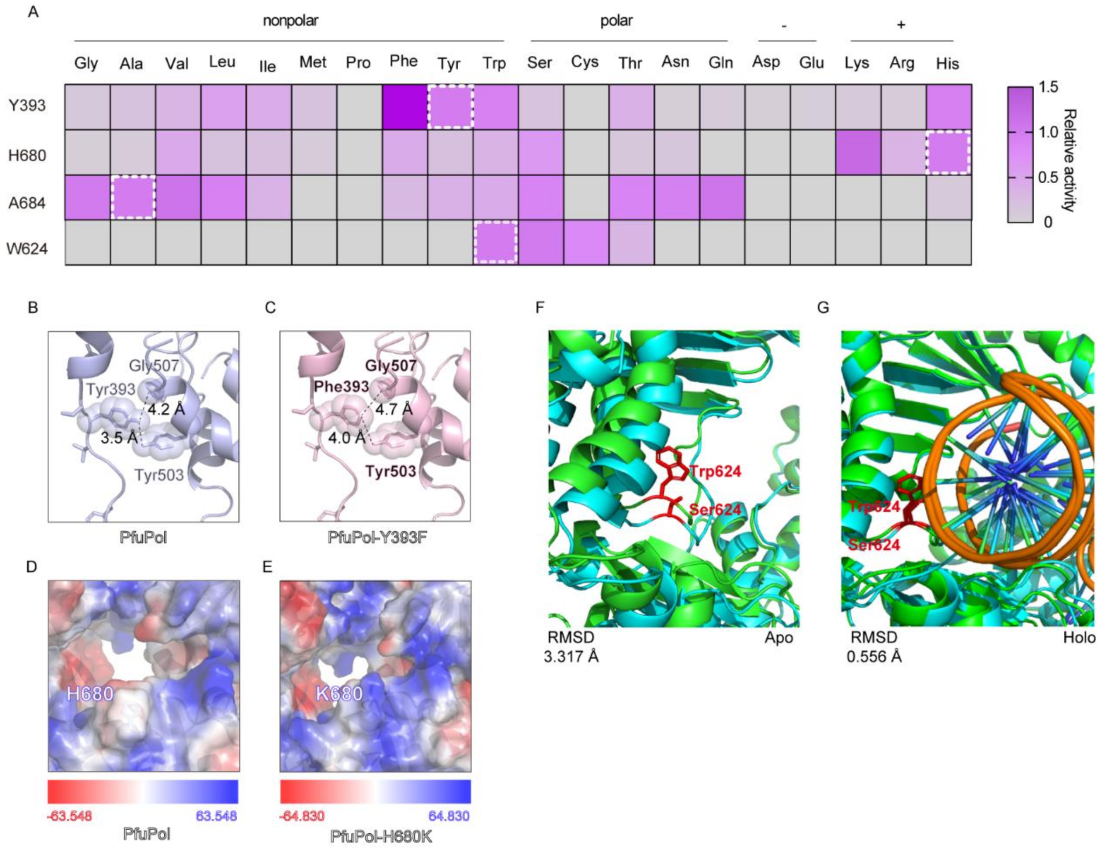


**Figure S7.** (**A**) The yield of the final PCR products was plotted as heatmap to represent the relative activity of PfuPol and its variants. The activity, corresponding to the yield of the final PCR products, is indicated by a color gradient from light gray (0) to dark purple (1.5). Wild-type PfuPol are labeled with a white dashed rectangle. (**B** and **C**) Structural illustration of the Y-GG/A motif of the wild-type PfuPol (**B**) and the variant PfuPol-Y393F (**C**). (**D** and **E**) Cross section view of the positions of the mutated residues in PfuPol. Wild-type PfuPol (**D**) and the variant PfuPol-H680K (**E**) is shown as an electrostatic potential surface (red, negative; blue, positive). (**F**) The structure of free PfuPol, shown in green, is aligned with the corresponding PfuPol-W624S structure displayed in lightblue. Sidechains are indicated in red. (**G**) The structure of PfuPol-DNA complex, shown in green, is aligned with the corresponding PfuPol-W624S-DNA complex structure displayed in lightblue. Sidechains are indicated in red. Structure figures were made using PyMOL.

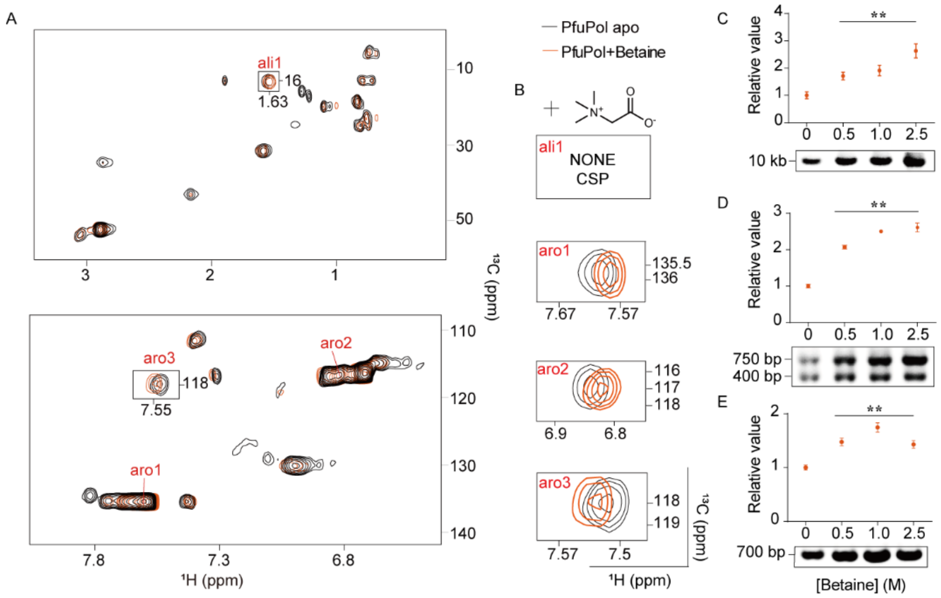


**Figure S8.** (**A**) 2D [$^{13}$C,$^{1}$H]-HMQC spectra of 50 μM [$^{13}$C]-labeled PfuPol acquired on the 600 MHz ($^{1}$H frequency) spectrometer at 298 K in the absence (black) and presence (orange) of 1M betaine. The spectrum was processed with a positive base level of 25000. (**B**) Enlarged view of NMR peaks of hotspot residues, which show CSPs upon adding 1 M betaine. The amplification of different DNA templates by PfuPol in the presence of different concentrations of betaine ranging from 0 M to 2.5 M. All PCR assays were performed as described in the methods. The PCR amplification with different templates: a 10 kb plasmid DNA template (**C**, with a target length of 10 kb), an Alpaca cDNA template (**D**, with target length of 750 bp and 400 bp) and a mouse tail gDNA template (**E**, with a target length of 700 bp). Intensities of gel bands were quantified and plotted against the concentration of betaine, showing on top of the corresponding gel image. Results expressed as mean ± SD of 3 independent biological replicates. Data were analyzed using unpaired Student's test, **P < 0.01.

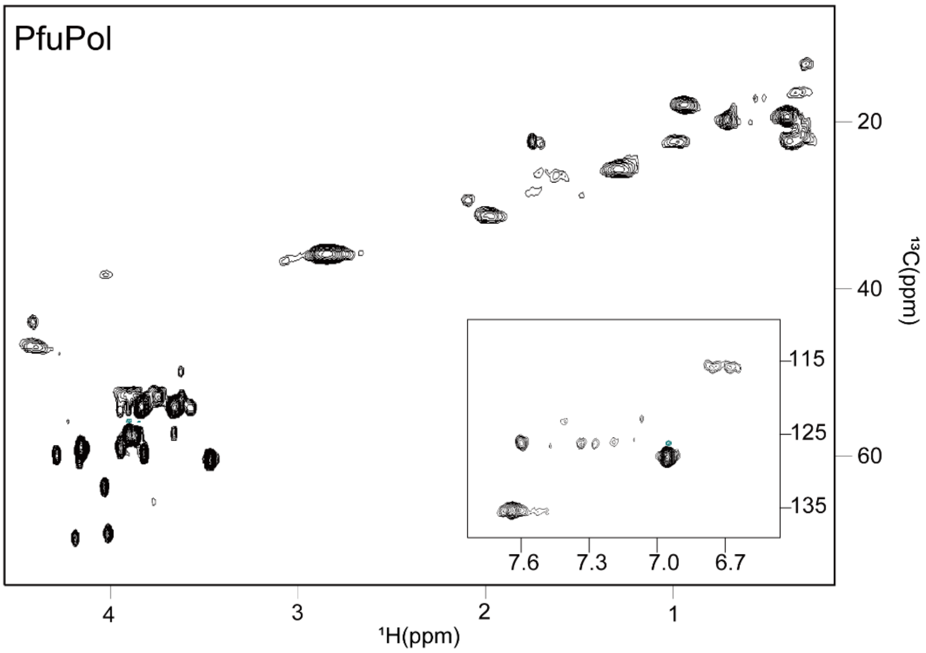


**Figure S9.** 2D [$^{13}$C,$^{1}$H]-HMQC spectra of 1 mM unlabeled PfuPol acquired on the 800 MHz ($^{1}$H frequency) spectrometer at 298 K.

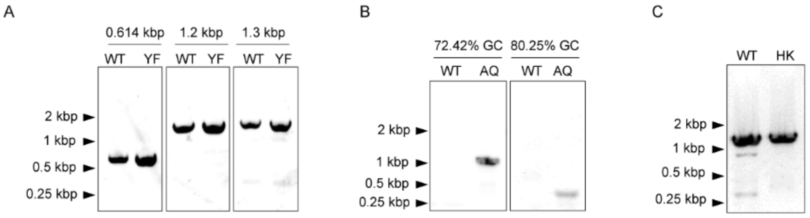


**Figure S10.** (**A**) PCR amplification of human gDNA substrate (with target length 614 bp, 1.2 kb and 1.3 kb). WT: wild-type PfuPol; YF: PfuPol-Y393F. (**B**) PCR amplification of rice cDNA substrate (with target length 471 bp, 80.25% GC, and 1.17 kb ,72.42% GC). WT: wild-type PfuPol; AQ: PfuPol-A684Q. (**C**) PCR amplification of cotton gDNA substrate (with target length 1.3 kb). WT: wild-type PfuPol; HK: PfuPol-H680K.

### Supplementary Tables

**Table S1.** Synthesized oligonucleotides.

| Name | Sequence |
|---|---|
| Hairpin DNA (T) | 5'-CCAGCATTAT GAAAGTGACA CGTGCACCAT TGGTGCACGT G-3' |
| Hairpin DNA (E) | 5'-CCAGCATTAT GAAAGTGACA CGTGCACCAT TGGTGCACGT GTCACTTTCA TAATGCTGG-3' |
| Y393N F | 5'-CGCCTGCGCG AATCCNNKAC CGGCGGTTTC GTG-3' |
| Y393N R | 5'-CACGAAACCG CCGGTMNNGG ATTCGCGCAG GCG-3' |
| H680N F | 5'-ATCACGCGTC CGCTGNNKGA ATACAAAGCC ATC-3' |
| H680N R | 5'-GATGGCTTTG TATTCMNNCA GCGGACGCGT GAT-3' |
| A684N F | 5'-CTGCACGAAT ACAAANNKA TCGGTCCGCAT GTT-3' |
| A684N R | 5'-AACATGCGGA CCGATMNNTT TGTATTCGTG CAG-3' |
| W624N F | 5'-AATTGTTCGC CGTGACNNKT CCGAAATCGC TAAAG-3 |
| W624N R | 5'-CTTTAGCGAT TTCGGAMNNG TCACGGCGAA CAATT-3 |
| Y393F F | 5'-CGCCTGCGCG AATCCTTTAC CGGCGGTTTC GTG-3' |
| Y393F R | 5'-CACGAAACCG CCGGTAAAGG ATTCGCGCAG GCG-3' |
| H680K F | 5'-ATCACGCGTC CGCTGAAAGA ATACAAAGCC ATC-3' |
| H680K R | 5'-GATGGCTTTG TATTCTTTCA GCGGACGCGT GAT-3' |
| A684Q F | 5'-CTGCACGAAT ACAAACAGA TCGGTCCGCAT GTT-3' |
| A684Q R | 5'-AACATGCGGA CCGATCTGTT TGTATTCGTG CAG-3' |
| W624S F | 5'-AAATTGTTCG CCGTGACAGC TCCGAAATCG CTAAAG-3' |
| W624S R | 5'-CTTTAGCGAT TTCGGAGCTG TCACGGCGAA CAATTT-3' |
| PP F | 5'-GCAGAAAAAG AAAAACTG-3' |
| PP R | 5'-GCGTTTCGCG CGCTGCAG-3' |
| Long F | 5'-ATCCTGGACG TGGATTACAT CACC-3' |
| Long R | 5'-CATATGACGA CCTTCGATAT GGCC-3' |
| Pucv F | 5'-GAATTCGAGC TCGGTACCC-3' |
| Pucv R | 5'-ACTGGCCGTC GTTTTAC-3' |
| LucB F | 5'-GGGGGATCCA TGAATTTTGG ATTATTCTTC-3' |
| LucB R | 5'-GGGAAGCTTT TACGAGCTTG GTAAATTC-3' |
| BCMA F | 5'-CTTGGATCCG GCACCAACGC GATTC-3' |
| BCMA Sub | 5'-CCAACGCGAT TCTGTGGACG AGCCTGGGCC TGAGCCTGAT TATTAGCCTG GCGGTGTTTG TGCTGATGTT TCTGCTGCGC AAAATTAACA GCTA-3' |
| BCMA R | 5'-GGGAAGCTTT TAGCTGTTAA TTTTG-3' |
| Os pro1 F | 5'-ATAACACATC CCTCCTTTTG TTATTTTATA TAC-3' |
| Os pro1 R | 5'-AGCTACAGCC TACACAGC-3' |
| Os pro2 F | 5'-GCCCTTTGGG ATCAAGCAAA TTAAACGC-3' |
| Os pro2 R | 5'-GGCTCCTGCT CTCTCCCC-3' |
| Os pro3 F | 5'-CTTCTATATA TAAAGTCATT TTCAACAATT ATATTAATTC C-3' |

| | |
|---|---|
| Os pro3 R | 5'-CTCTTCCTCT TGCTGTTCTT GCTTCTTC-3' |
| Os CDS1 F | 5'-ATGGCGTCCT ACGACAAGGC C-3' |
| Os CDS1 R | 5'-TTATTTGCTT GCATCTTGGG GATCTTGATT CTC-3' |
| Gh gDNA F | 5'-GGTACCGAGA TACAGAAGTT TTAG-3' |
| Gh gDNA R | 5'-GTCGACCATC TAGAACAGGA GTGAC-3' |
| Ap cDNA F | 5'-GTCCTGGCTC TCTTCTACAA GG-3' |
| Ap cDNA R | 5'-GGTACGTGCT GTTGAACTGT TCC-3' |
| Mt gDNA F | 5'-AACCCAGAAG ACAGGTGGAA AG-3' |
| Mt gDNA R | 5'-GCCAGATTAC GTATATCCTG GCAG-3' |

**Table S2.** Summary of model quality scores for complex components.

| Complex Component | Hairpin DNA | Nb2 | Nb4 |
|---|---|---|---|
| PTM | 0.92 | 0.89 | 0.87 |
| iPTM | 0.72 | 0.85 | 0.85 |
| Haddock score | - | –63.6 ~ –110.99 | –86.81 ~ –124.10 |

**Table S3.** Summary of hotspots assignment.

| hotspot | | Potential residue | | Distance to residue (Å) | | | | | |
|---|---|---|---|---|---|---|---|---|---|
| Name | Pattern* | Position | P(%) of type | hairpin DNA | | Nb2 | | Nb4 | |
| ali1 | S,S,S | A257 | 78.8 | 21.0 ± 7.1 | ❌ | 7.7 ± 1.9 | ✅ | 11.4 ± 1.9 | ❌ |
| **ali1** | S,S,S | **A684** | **78.8** | 4.8 ± 0.4 | ✅ | 4.6 ± 0.6 | ✅ | 4.2 ± 0.6 | ✅ |
| ali1 | S,S,S | M255 | 21.2 | 10.0 ± 4.9 | ❌ | 4.2 ± 0.6 | ✅ | 4.4 ± 0.8 | ✅ |
| aro1 | S,S,W | H270 | 100 | 8.3 ± 2.7 | ❌ | 5.4 ± 1.3 | ✅ | 7.4 ± 2.1 | ❌ |
| **aro1** | S,S,W | **H680** | **100** | 6.7 ± 1.0 | ✅ | 6.4 ± 1.1 | ✅ | 10.8 ±2.2 | ✅ |
| aro1 | S,S,W | H688 | 100 | 5.3 ± 1.0 | ✅ | 5.1 ± 1.6 | ✅ | 5.2 ± 1.5 | ❌ |
| aro2 | S,W,S | Y154 | 70.3 | 30.0 ± 0.0 | ❌ | 4.9 ± 1.8 | ❌ | 4.8 ± 1.8 | ✅ |
| aro2 | S,W,S | Y226 | 70.3 | 25.0 ± 7.1 | ❌ | 7.8 ± 2.0 | ✅ | 6.0 ± 2.6 | ✅ |
| aro2 | S,W,S | Y269 | 70.3 | 10.0 ± 2.2 | ❌ | 4.2 ± 0.6 | ❌ | 4.0 ± 0.0 | ✅ |
| aro2 | S,W,S | Y281 | 70.3 | 8.0 ± 0.0 | ❌ | 4.4 ± 1.3 | ❌ | 4.0 ± 0.0 | ✅ |
| aro2 | S,W,S | Y385 | 70.3 | 7.0 ± 1.7 | ✅ | 4.8 ± 2.5 | ❌ | 5.6 ± 2.4 | ✅ |
| **aro2** | S,W,S | **Y393** | **70.3** | 4.2 ± 0.5 | ✅ | 10.6 ± 2.1 | ✅ | 6.6 ± 1.1 | ✅ |
| aro2 | S,W,S | Y508 | 70.3 | 4.0 ± 0.0 | ✅ | 6.4 ± 2.6 | ❌ | 6.2 ± 2.4 | ✅ |
| aro2 | S,W,S | Y672 | 70.3 | 5.2 ± 1.0 | ✅ | 9.8 ± 2.4 | ✅ | 8.4 ± 1.3 | ❌ |
| aro2 | S,W,S | Y682 | 70.3 | 4.0 ± 0.0 | ✅ | 5.3 ± 2.5 | ❌ | 4.2 ± 0.6 | ✅ |
| aro2 | S,W,S | Y710 | 70.3 | 7.2 ± 1.1 | ✅ | 6.9 ± 1.8 | ❌ | 6.5 ± 0.9 | ✅ |
| aro2 | S,W,S | Y740 | 70.3 | 4.0 ± 0.0 | ✅ | 8.0 ± 2.3 | ✅ | 10.8 ± 2.1 | ❌ |
| aro2 | S,W,S | W624 | 14.9 | 4.5 ± 0.7 | ✅ | 5.8 ± 1.2 | ❌ | 7.0 ± 0.9 | ✅ |
| aro2 | S,W,S | H270 | 4.4 | 8.3 ± 2.7 | ❌ | 5.4 ± 1.3 | ❌ | 7.4 ± 2.1 | ✅ |
| aro2 | S,W,S | H680 | 4.4 | 6.7 ± 1.0 | ✅ | 6.4 ± 1.1 | ❌ | 10.8 ±2.2 | ❌ |
| aro2 | S,W,S | H688 | 4.4 | 5.3 ± 1.0 | ✅ | 5.1 ± 1.6 | ❌ | 5.2 ± 1.5 | ✅ |
| **aro3** | **S,S,S** | **W624** | **85.1** | 4.5 ± 0.7 | ✅ | 5.8 ± 1.2 | ✅ | 7.0 ± 0.9 | ✅ |
| aro3 | S,S,S | H270 | 14.9 | 8.3 ± 2.7 | ❌ | 5.4 ± 1.3 | ✅ | 7.4 ± 2.1 | ✅ |
| aro3 | S,S,S | H680 | 14.9 | 6.7 ± 1.0 | ✅ | 6.4 ± 1.1 | ✅ | 10.8 ±2.2 | ❌ |
| aro3 | S,S,S | H688 | 14.9 | 5.3 ± 1.0 | ✅ | 5.1 ± 1.6 | ✅ | 5.2 ± 1.5 | ✅ |

* S: Significant CSP; W: Non-significant CSP.
e.g. S,S,W: Significant CSP with hairpin DNA, Significant CSP with Nb2, Non-significant CSP with Nb4.

**Table S4.** Summary of the steady-state kinetics analyses of PfuPol and its variants.

| Enzyme | $k_{cat}$ ($s^{-1}$) | $K_M$ (μM) | $k_{cat}$ / $K_M$ ($M^{-1}$ $s^{-1}$) | Error rate (× $10^{-6}$) |
|---|---|---|---|---|
| PfuPol | 190.70 ± 11.07 | 0.45 ± 0.07 | (4.42 ± 0.70) × $10^8$ | 3.80 ± 0.37 |
| PfuPol-Y393F | 258.00 ± 8.22 | 0.43 ± 0.08 | (6.00 ± 1.13) × $10^8$ | 2.68 ± 0.28 |
| PfuPol-H680K | 173.28 ± 5.36 | 0.31 ± 0.06 | (5.59 ± 1.10) × $10^8$ | 3.24 ± 0.46 |
| PfuPol-A684Q | 194.90 ± 2.36 | 0.34 ± 0.03 | (5.73 ± 0.51) × $10^8$ | 3.99 ± 0.46 |
| PfuPol-W624S | 255.80 ± 12.64 | 0.46 ± 0.06 | (5.56 ± 0.78) × $10^8$ | 3.87 ± 0.32 |

**Table S5.** Summary of the steady-state kinetics analyses of PfuPol and its variants.

| Enzyme | $k_{cat}$ ($s^{-1}$) | $K_M$ (μM) | $k_{cat}/K_M$ ($M^{-1}s^{-1}$) |
|---|---|---|---|
| PfuPol-Y393F/H680K | 1571.7 ± 139.3 | 0.33 ± 0.07 | (4.76 ± 1.10) × $10^9$ |
| PfuPol-Y393F/A684Q | 378.1 ± 31.3 | 0.42 ± 0.09 | (0.90 ± 0.20) × $10^9$ |
| PfuPol-H680K/A684Q | 1341.2 ± 77.2 | 0.29 ± 0.05 | (4.62 ± 0.84) × $10^9$ |
| PfuPol-Y393F/H680K/A684Q | 292.9 ± 16.0 | 0.35 ± 0.04 | (0.84 ± 0.11) × $10^9$ |
| PfuPol-Y393F/W624S/A684Q | 230.6 ± 10.9 | 0.57 ± 0.07 | (0.41 ± 0.05) × $10^9$ |
